# TLR4^+^ Dermal fibroblasts induce acute and transitional pain states

**DOI:** 10.1101/2025.03.07.641865

**Authors:** Melissa E. Lenert, Nilesh M. Agalave, Emily K. Debner, Jessica A. Tierney, Syed A. Naqvi, Andreas M. Chavez, Marilyn Dockum, Phil Albrecht, Frank L. Rice, Theodore J. Price, Erica L. Sanchez, Michael D. Burton

## Abstract

The prominence of non-neuronal cells driving pain states has gained attention in recent years. Fibroblasts, a major stromal cell, perform essential functions during inflammation, tissue remodeling, and wound healing; however, recent studies suggest that fibroblasts may play a role in pain. Toll-like receptor 4 (TLR4) is an essential component of the innate immune system and activation of the receptor promotes pain. This study utilized a novel mouse model with dermal fibroblast specific expression of TLR4 on a TLR4-null background, which allows us to understand the sufficiency of skin fibroblast activation in pain development. Here we demonstrate that dermal fibroblast activation induces both acute inflammatory pain and hyperalgesic priming in both male and female mice. *In vivo*, activated dermal fibroblasts change cellular morphology in mice and humans. *In vitro* we observed pro-inflammatory cytokine production and activation of calcium signaling pathways. These data demonstrate that dermal fibroblast activation can cause acute pain and drive mechanisms involved in the transition to chronic pain.

**Summary:** Non-neuronal cells are an emerging target for the development of therapeutics for chronic pain. Activation of dermal fibroblasts via TLR4 is sufficient to induce inflammatory pain and hyperalgesic priming in male and female mice.

## Introduction

Over 100 million Americans experience chronic pain and chronic pain-induced disability and current therapeutics are lacking in efficacy and safety (Vos et al., 2015). Important therapeutic strategies should target early pain and the acute-to-chronic pain transition. Moreover, neuroimmune interactions may explain current deficits in pain treatment (Lenert et al., 2021; Pinho-Ribeiro et al., 2017; Szabo-Pardi et al., 2021a), but there is still a pressing need to identify novel targets for analgesic development. In particular, the prominent role of non-neuronal cells driving pain states has gained much attention in recent years (Grace et al., 2021). In this study, we utilized a novel mouse model to examine the role of dermal fibroblast-TLR4 activation in inflammatory pain and acute-to-chronic priming. A few studies have begun to reveal the important role of various fibroblast populations in pain processing (Singhmar et al., 2020; Wei et al., 2014); however, ours is the first study to show that direct dermal fibroblast activation alone is sufficient to cause both acute pain sensitivity and hyperalgesic priming.

Fibroblasts are a heterogeneous population of mesenchymal stromal cells traditionally recognized for synthesizing and remodeling extracellular matrix (ECM) during wound healing, tissue remodeling, inflammation, and several other processes (Ascensión et al., 2021; Deng et al., 2021; Lynch and Watt, 2018). During inflammatory conditions or fibrosis, fibroblasts produce pro-inflammatory cytokines (IL-6, IL-1β) and have been shown to modulate the extracellular environment to facilitate immune cell recruitment (Laurent et al., 2007; Tse et al., 2014; Wei et al., 2014). Dermal fibroblasts produce collagen and other ECM components that provide structural integrity to the skin (Laurent et al., 2007; Tracy et al., 2016). Dermal fibroblasts also play a major role in the immune function of the skin by providing structural defense, producing cytokines and chemokines, and even directly responding to pathogens via expression of pattern recognition receptors (Deng et al., 2021; Nguyen and Soulika, 2019).

TLR4 is a pattern-recognition receptor (PRR) expressed on many cell types, including fibroblasts, that orchestrates the inflammatory response to injury and infection as well as pain development in response to both pathogens and tissue damage (Bruno et al., 2018; Sauer et al., 2014). In the periphery, TLR4 signaling mediates cell-specific responses to contribute to acute and chronic pain states (Burton et al., 2019; Rudjito et al., 2021; Szabo-Pardi et al., 2021a). Recent evidence has shown that dermal fibroblasts use a TLR4-dependent mechanism to respond to inflammation and/or injury (Bhattacharyya et al., 2018; Liu et al., 2018); however, the mechanism by which fibroblasts contribute to pain states via TLR4 signaling is not understood.

In the current study, we used a newly developed and validated genetic model to reactivate TLR4 on dermal Fibroblast-Specific Protein 1 (FSP1)^+^ fibroblasts on a whole-body TLR4-null background. We characterized FSP1cre expression in dermal fibroblasts, *in vivo*, using two-photon imaging. Using LPS-FITC (lipopolysaccharide, TLR4 agonist), we observed the FITC-tagged LPS bound to tdTomato (FSP1cre^+^) positive fibroblasts in the hindpaw of alive animals. Dermal fibroblast-TLR4 activation alone was sufficient to cause mechanical hypersensitivity after a single intraplantar injection of LPS in both male and female mice, similarly to their wild-type counterparts. Furthermore, activation of fibroblast TLR4 is sufficient to drive hyperalgesic priming. We also conducted, *in vivo* morphological changes in dermal fibroblasts following activation in mouse and human skin. Finally, we utilized *in vitro* activation of dermal fibroblasts to understand intracellular signaling mechanisms that occur in fibroblasts following TLR4 activation. Taken together, this study provides novel evidence that fibroblast activation alone is sufficient to cause inflammatory pain via TLR4 activation and are involved in the transition to chronic pain through neuronal hyperalgesic priming.

## Results

### *In vivo,* dermal FSP1^+^ fibroblasts are responsive to the TLR4 agonist (LPS)

We utilized our previously described reporter mouse [FSP1^tdt^: FSP1cre^+^ fibroblasts express tdTomato and have wild-type (WT) expression of TLR4; **Figure 1A-B**; (Szabo-Pardi et al., 2019)] to understand the expression pattern of FSP1^+^ fibroblasts in the skin of the plantar surface of the hind paw. The presence of FSP1cre-dependent tdTomato expression can be visualized in live mice prior to experiments by using green LED flashlight with red filter glasses. In the presence of FSP1cre, tdTomato is expressed robustly in the skin and can be visualized easily in the ears (**Figure 1B**) in FSP1cre(^+^) mice (right) compared to their Cre-negative littermates (left).

**Figure 1.**
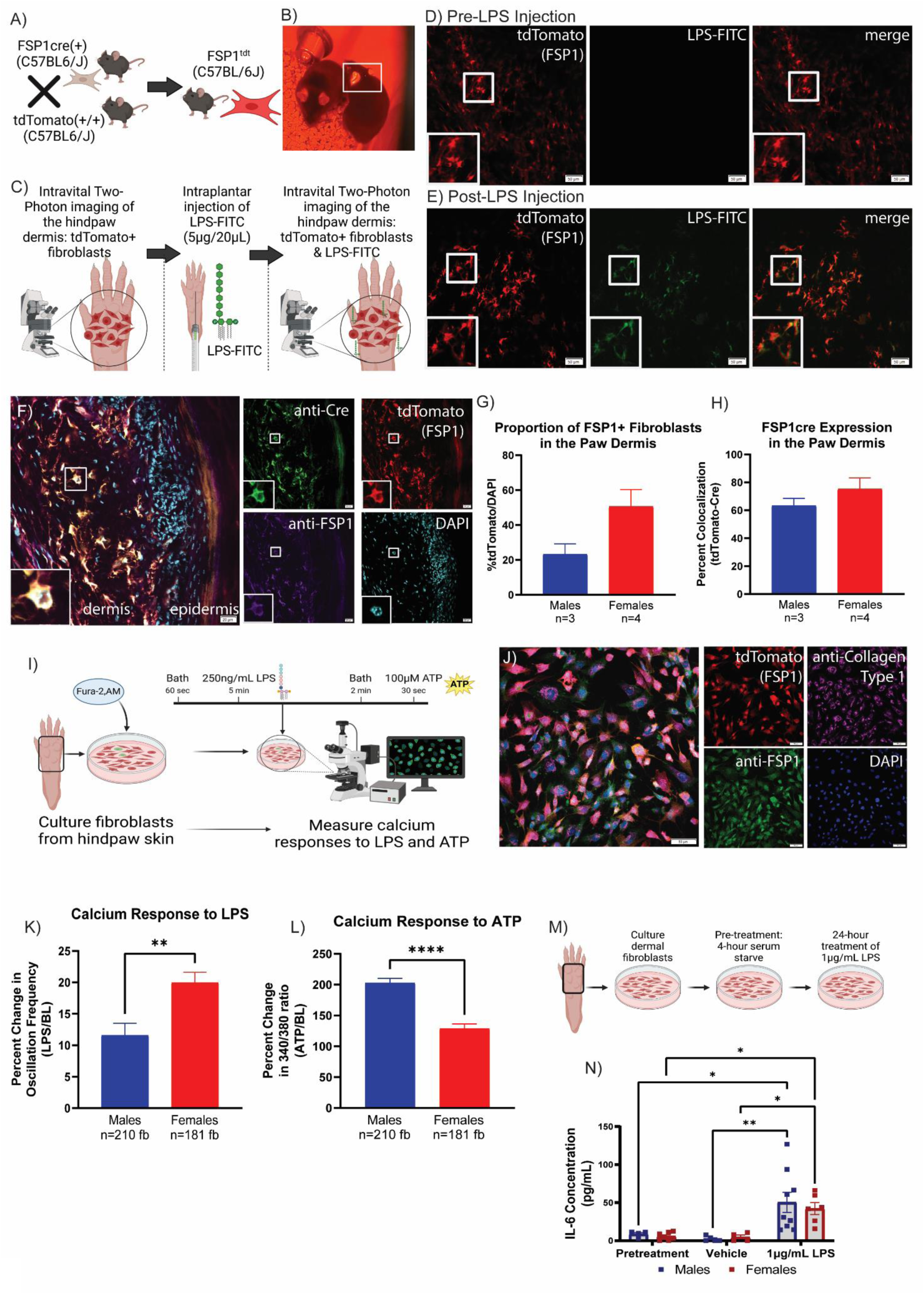
FSP1^+^ dermal fibroblasts respond to the TLR4 agonist LPS. (A) Generation of FSP1^tdt^ reporter mice. (B) Visualization of tdTomato expression in the skin of FSP1cre (^+^) (right) and FSP1cre (-) (left) mice. (C) Graphic depicting the experimental schematic for intravital two-photon imaging of LPS-FITC uptake by dermal fibroblasts expressing tdTomato. (D) FSP1^tdt^ hind paw dermis prior to intraplantar injection of LPS-FITC. Scale bar = 50µm. (E) FSP1^tdt^ hind paw dermis 1h after intraplantar injection of LPS-FITC. Scale bar = 50µm. (F) Representative confocal microscopy of Cre recombinase (green), FSP1 (purple) and tdTomato (red) immunohistochemistry in the hind paw skin from FSP1^tdt^ mice. Scale bar = 20µm. (G) Quantification of the percentage of tdTomato (^+^) cells in the hind paw dermis from FSP1^tdt^ mice compared to the total number of DAPI ^+^ cells. (Males: n=3 mice, Females: n=4 mice). (H) Quantification of FSP1cre expression via colocalization of tdTomato, Cre recombinase, and FSP1 in the hindpaw. (Males: n=3 mice, Females, n=4 mice). (I) Graphic depicting the use of fura-2 AM for imaging cultured dermal fibroblast calcium responses to LPS and ATP. (J) Representative confocal microscopy of FSP1 (green), Collagen Type 1 (purple) and tdTomato (red) immunocytochemistry of cultured dermal fibroblasts from the hind paw skin of FSP1^tdt^ mice. Scale bar = 50µm. (K) Representative calcium traces of five randomly selected fibroblasts during baseline (0- 60s), 250 ng/mL LPS treatment (60-360s), wash-out (360-480s), and 100µM ATP (480- 500s). FB = fibroblast. (L) Representative calcium traces zoomed in on oscillations observed during baseline and LPS treatment. (M) Percent change in fibroblast oscillation frequency after treatment with 250ng LPS (Males: n=210 fb, n=2 mice; Females: n=181 fb, n=2 mice). (N) Percent change in Fura 2-AM 340/380 ratio after treatment with 100µM ATP (Males: n=210 fb, n=2 mice; Females: n=181 fb, n=2 mice). (O) Experimental timeline for treatment of cultured dermal fibroblasts with LPS. (P) IL-6 concentration in media collected from cultured fibroblasts treated with 1µg/mL LPS for 24h. \**p*<0.05, \*\**p*<0.01, \*\*\*\**p*<0.0001 by unpaired two-tailed t-test (G, H, M, N) or Ordinary Two-Way ANOVA (P).

Using intravital two-photon imaging, we observed the uptake of LPS-FITC into tdTomato (^+^) fibroblasts in the hind paw dermis to confirm that dermal FSP1cre+ fibroblasts express functional TLR4 (**Figure 1C**). Prior to LPS-FITC injection, expression of tdTomato can be observed in dermal fibroblasts with little to no autofluorescence on the FITC channel (**Figure 1D**). After an intraplantar injection of LPS-FITC, co-localization of tdTomato (FSP1) and FITC (LPS) can be observed throughout the hind paw dermis indicating uptake of LPS into dermal fibroblasts (**Figure 1E**).

We sought to assess the expression pattern of FSP1cre in the hind paw dermis using our FSP1^tdt^ mice. We utilized immunohistochemistry to determine the expression of FSP1cre in the hind paw by staining for FSP1 and Cre recombinase in our FSP1^tdt^ mice (**Figure 1F**). We measured the percentage of tdTomato (^+^) cells out of the total cell count (via DAPI) and found that FSP1cre (^+^) fibroblasts (tdTomato-expressing cells) make up 23.17% (± 6.05%) and 50.56% (± 9.78%) of total live cells in the hind paw dermis in males and females, respectively (**Figure 1G**). Further, tdTomato and Cre recombinase expression is observed in 63.3% (± 5.3%) and 75.2% (± 7.98%) of FSP1- expressing cells in the dermis in males and females, respectively (**Figure 1H**). No statistically significant differences were observed in FSP1cre expression between males and females.

Finally, we performed a morphology analysis using Imaris software to determine the 3D shape of dermal fibroblasts from naïve FSP1^tdt^ mice (**Supplemental Figure 1A**). The cell volume of dermal fibroblasts from naïve FSP1^tdt^ males (1176 ± 42.4 µm^3^) are not significantly different than those from naïve FSP1^tdt^ females (937.9 ± 240.2 µm^3^) (**Supplemental Figure 1B**). The oblate ellipticity (flattening) of dermal fibroblasts from naïve FSP1^tdt^ males (0.375 ± 0.004 AU) is not significantly different from naïve FSP1^tdt^ females (0.405 ± 0.026 AU) (**Supplemental Figure 1C**). The prolate ellipticity (elongation) of dermal fibroblasts from naïve FSP1^tdt^ males (0.452 ± 0.005 AU) is not significantly different from naïve FSP1^tdt^ females (0.421 ± 0.021 AU) (**Supplemental Figure 1D**).

### *In vitro*, primary cultured fibroblasts from the hind paw skin respond to TLR4 agonist (LPS)

To determine the mechanisms by which dermal fibroblasts respond to LPS, we isolated and cultured fibroblasts from the hind paw skin of male and female WT mice (**Figure 1I**). We first verified that the cultured cells were fibroblasts by staining cultured fibroblasts from FSP1^tdt^ mice for FSP1 and Type 1 Collagen, both widespread markers of stromal fibroblasts (**Figure 1J**).

Next, we performed calcium imaging of cultured fibroblasts from WT male and female mice to determine whether agonism of TLR4 via LPS would affect calcium oscillations when measured using fura-2 AM, which would target all the fibroblasts in the dish regardless of FSP1 expression . Representative calcium trace (**Supplemental Figure 1E**) compared to our calcium oscillation analysis pipeline output (**Supplemental Figure 1F**) shows identification of prominent peaks and oscillation frequency. Fibroblasts were treated with LPS and then ATP was applied as a positive control after a 2-minute washout period **(Supplemental Figure 1G)**. At baseline, male dermal fibroblasts have an oscillation frequency of 14.31±0.19 oscillations per minute (osc/min) whereas female dermal fibroblasts have an oscillation frequency of 11.98±0.17 osc/min (**Supplemental Figure 1H**). After application of 250ng/mL LPS, calcium oscillation frequency increased by 11.59±1.93% in male fibroblasts (n=210 fibroblasts; n=2 mice) and by 19.97±1.65% in female fibroblasts (n=181 fibroblasts; n=2 mice), with a significantly greater increase in females compared to males (**Figure 1K**, t (378) = 3.242, *p*=0.0013, unpaired two-tailed t-test). Following LPS application and a 2-minute washout period, application of 100µM ATP (positive control) caused a sharp calcium influx in both male (203±7.4% increase in fluorescence intensity) and female (128.6±7.8% increase in fluorescence intensity) fibroblasts, with a significantly greater response to ATP in males compared to females (**Figure 1L**, t (390) = 6.910, *p*<0.0001, unpaired two-tailed t-test).

Additionally, we measured calcium oscillations with 1µg/mL LPS (**Supplemental Figure 1I**). After application of 1µg/mL LPS, calcium oscillation frequency increased by 5.04±1.31% in male fibroblasts (n=221 fibroblasts; n=3 mice) and by 14.1±1.9% in female fibroblasts (n=142 fibroblasts; n=3 mice), with a significantly greater increase in females compared to males (**Supplemental Figure 1J**, t (361) = 4.058, *p*<0.0001, unpaired two-tailed t-test). Following LPS application and a 2-minute washout period, application of 100µM ATP caused a sharp calcium influx in both male (242.1±7.85% increase in fluorescence intensity) and female (262.0±10.9% increase in fluorescence intensity) fibroblasts, with no difference between males and females (**Supplemental Figure 1K**).

Finally, we measured IL-6 production by cultured fibroblasts treated with 1µg LPS for 24 hours after a 4-hour serum starve (**Figure 1M, N**). Treatment with LPS significantly increased IL-6 production in both WT male and female fibroblasts compared to both pre-treatment groups and vehicle-treated groups (LPS treatment: F (2, 29) = 13.42, *p*<0.0001, Two-Way ANOVA) with no sex differences observed.

Further, to assess calcium oscillation frequency in specifically FSP1^+^ fibroblasts, FSPcre mice were crossed with mice expressing GCaMP6f – an intracellular calcium sensor (FSP^Salsa^, **Supplemental Figure 2A**). Extracellular matrix can affect cellular function and activity, so we assessed calcium oscillations from fibroblasts cultured on poly-D lysine coated or poly-D lysine and collagen coated plates. Similar to WT cultures, after application of 1µg/mL LPS, female FSP1^+^ dermal fibroblasts have increased calcium oscillations compared to male FSP1+ dermal fibroblasts, regardless of plate coating while application of 100μM ATP produced larger calcium responses in male fibroblasts compared to female fibroblasts (**Supplemental Figure 2B-G**).

### Generation and validation of FSP1^TLR4LoxTB^ mouse line

To assess whether activation of dermal fibroblasts via TLR4 signaling was sufficient to induce pain responses, we crossed FSP1cre mice with TLR4^LoxTB^ mice (whole-body TLR4 null) to generate mice with fibroblast-specific expression of TLR4 (FSP1^TLR4LoxTB^) in the presence of FSP1cre (**Figure 2A**). Cre-negative littermate controls have little to no expression of TLR4 (TLR4^LoxTB^) (Jia et al., 2021; Szabo-Pardi et al., 2021a). PCR and gel electrophoresis were used to confirm genotypes (**Figure 2B, C**).

**Figure 2.**
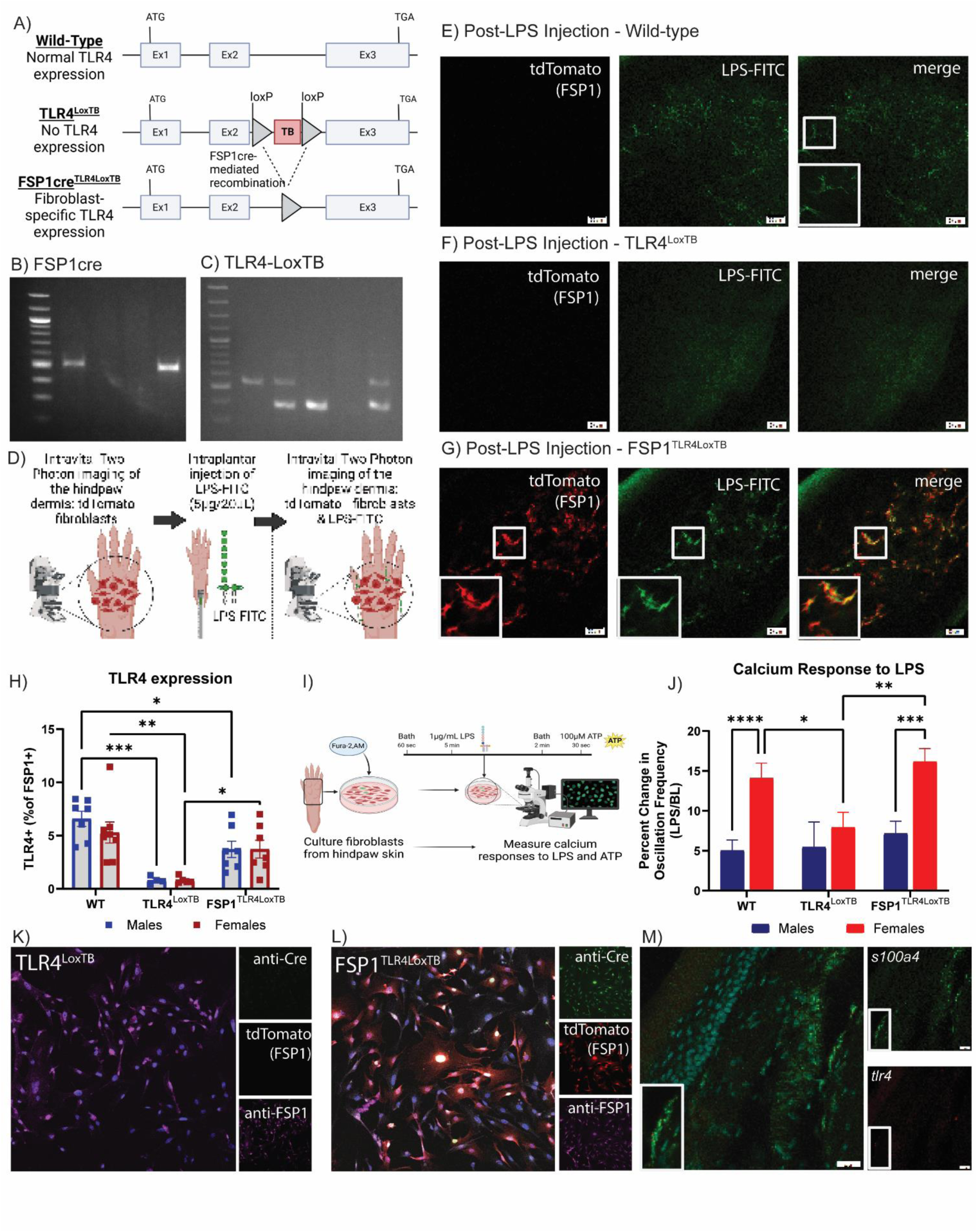
Generation and Validation of FSP1^TLR4LoxTB^ mice. (A) Generation of TLR4^LoxTB^ and FSP1^TLR4LoxTB^ mice for fibroblast-specific expression of TLR4. (B) Representative PCR and gel electrophoresis of FSP1cre (FSP1cre = 500bp). (C) Representative PCR and gel electrophoresis of TLR4-LoxTB (TLR4 WT = 523bp; TLR4-LoxTB = 370bp). (D) Graphic depicting the experimental schematic for intravital two-photon imaging of LPS-FITC uptake by dermal fibroblasts expressing tdTomato. (E-G) Wild-type (E), TLR4^LoxTB^ (F), and FSP1^TLR4LoxTB^ (G) hind paw dermis 1h after intraplantar injection of LPS-FITC. Scale bar = 50µm. (H) TLR4 protein expression in FSP1^+^ dermal fibroblasts assessed via flow cytometry. (I) Graphic depicting the use of fura-2 AM for imaging cultured dermal fibroblast calcium responses to LPS and ATP. (J) Percent change in calcium oscillation frequency after treatment with 1µg/mL LPS. (K-L) Representative confocal microscopy of Cre Recombinase (green), tdTomato (red) and FSP1 (purple) immunocytochemistry of cultured dermal fibroblasts from TLR4^LoxTB^ (H) and FSP1^TLR4LoxTB^ (I) mice. Scale bar = 50µM. (J) Representative confocal microscopy of RNA *in situ* hybridization of *s100A4* (green) and *tlr4* (red) mRNA expression in the hind paw dermis of wild-type mice. Cre recombinase expression fibroblasts cultured from in TLR4^LoxTB^ (K) and FSP1^TLR4LoxTB^ (L). (M) *in situ* hybridization of WT mouse hind paw dermis targeting tlr4 (red) and s100a4 (green) mRNA. Scale bar = 20µM. \**p*<0.05, \*\**p*<0.01, \*\*\**p*<0.001, \*\*\*\**p*<0.0001 by Ordinary Two-Way ANOVA (H, J).

As with the FSP1^tdt^ mice, we utilized intravital two-photon imaging to assess the uptake of LPS-FITC in the hind paw dermis of WT, TLR4^LoxTB^, and FSP1^TLR4LoxTB^ mice and confirm the presence or absence of functional TLR4 in FSP1^TLR4LoxTB^ mice and TLR4^LoxTB^ mice, respectively (**Figure 2D**). Prior to LPS injection, tdTomato(^+^) fibroblasts can be observed in FSP1^TLR4LoxTB^ mice, but not TLR4^LoxTB^ and WT mice. In all three groups, little to no autofluorescence on the FITC channel is observed prior to LPS-FITC injection (**Supplemental Figure 3A-C**). In WT mice, the uptake of LPS-FITC can be observed in several cell types including those with fibroblast-like morphology (**Figure 2E**). In contrast, TLR4^LoxTB^ mice show no uptake of LPS-FITC and instead have diffusion of LPS-FITC throughout the dermis (**Figure 2F**). FSP1^TLR4LoxTB^ mice, which have FSP1^+^ fibroblast-specific expression of TLR4, show co-localization of LPS-FITC and tdTomato indicating uptake of LPS-FITC in FSP1cre (^+^) fibroblasts and successful re-expression and functionality of TLR4 in these cells (**Figure 2G**).

We used flow cytometry to measure the protein expression of TLR4 on FSP1^+^ fibroblasts from the hind paw skin of WT, TLR4^LoxTB^, and FSP1^TLR4LoxTB^ male and female mice (**Figure 2H**; **Supplemental Figure 4**). We identified low levels of TLR4 protein in plantar skin FSP1^+^ fibroblasts of WT male and female mice and successful re-expression of TLR4 in FSP1^TLR4LoxTB^ mice (genotype: F(2, 32) = 52.10, *p*<0.0001, Two-Way ANOVA). Our whole-body null TLR4^LoxTB^ animals express minimal TLR4 in fibroblasts populations, similar to our previous observations in neuronal and immune cell populations within the DRG (Szabo-Pardi et al., 2021a).

Next, we performed calcium imaging of cultured fibroblasts from male and female mice to determine whether re-expression of TLR4 in FSP1^+^ fibroblasts only would recapitulate the phenotype observed in WT fibroblasts (**Figure 2I-J**). As such, the WT fibroblast data shown in **Figure 2J** is the same as that shown in **Supplemental Figure 1I**. After application of 1µg/mL LPS, calcium oscillation frequency increased by 5.04±1.31% in male WT fibroblasts (n=221 fibroblasts; n=3 mice), by 5.45±3.15% in male TLR4^LoxTB^ fibroblasts (n=64 fibroblasts, n=3 mice), and by 7.17±1.54% in male FSP1^TLR4LoxTB^ fibroblasts (n=204 fibroblasts, n=4 mice). After application of 1µg/mL LPS, calcium oscillation frequency increased by 14.1±1.9% in female WT fibroblasts (n=142 fibroblasts; n=3 mice), by 7.93±1.9% in female TLR4^LoxTB^ fibroblasts (n=135 fibroblasts, n=3 mice), and by 16.16±1.6% in female FSP1^TLR4LoxTB^ fibroblasts (n=135 fibroblasts, n=4 mice). Independent of genotype, female fibroblasts had a significantly greater increase in calcium oscillation frequency in response to LPS application compared to male fibroblasts (sex: F(1, 895) = 19.73, *p*<0.0001, Two-Way ANOVA, **Figure 2J**). There was also a significant genotype effect (F(2, 895) = 3.104, p=0.454, Two-Way ANOVA, **Figure 2J**); However, no significant *post hoc* genotype differences were observed in male fibroblasts. In female fibroblasts, TLR4^LoxTB^ had a lower response to LPS application than both WT and FSP1^TLR4LoxTB^ fibroblasts. WT and FSP1^TLR4LoxTB^ fibroblasts have similar calcium responses to LPS.

In cultured dermal fibroblasts, no Cre recombinase or tdTomato expression is observed in TLR4^LoxTB^ mice (**Figure 2K**). In contrast, Cre recombinase expression can be observed in tdTomato (^+^) fibroblasts in FSP1^TLR4LoxTB^ mice (**Figure 2L**). Similar expression of FSP1 is observed in both TLR4^LoxTB^ and FSP1^TLR4LoxTB^ fibroblasts.

We used RNA *in situ* hybridization (ISH) to qualitatively confirm expression of TLR4 within FSP1^+^ fibroblasts within the hind paw dermis of WT mice (**Figure 2M**).

Expression of *S100A4* (FSP1) mRNA can be observed throughout the hind paw dermis. Expression of *tlr4* mRNA can also be observed throughout the dermis with low expression in fibroblasts and other cell types. We also performed RNA ISH on WT DRGs for expression of *Tlr4* and *S100A4* mRNA (**Supplemental Figure 5**). As expected, *Tlr4* mRNA can be observed in approximately 30% of sensory neurons within the DRG in addition to expression in non-neuronal cells (Jia et al., 2021). Little to no expression of *S100A4* was seen, indicating that FSP1^+^ fibroblasts are not localized within the DRG.

### Intraplantar injection of LPS induces pain-like behavior in both male and female mice

To investigate functionality of TLR4 on fibroblasts and assess the sufficiency of fibroblast activation on acute pain development and resolution, we measured mechanical sensitivity and facial grimacing following a single intraplantar of LPS (**Figure 3A**). There were no differences in mechanical sensitivity at baseline regardless of the presence of TLR4 in either males or females. After intraplantar injection of LPS, both male and female FSP1^TLR4LoxTB^ mice develop sensitivity similarly to WT mice, which peaks at 4-6h post injection and resolves at 24h post injection (**Figure 3B, C**). As expected, TLR4^LoxTB^ mice had no behavioral response to LPS injection thus confirming their TLR4-null phenotype. Though WT male mice have greater mechanical sensitivity than FSP1^TLR4LoxTB^ males, there were not *post hoc* differences observed between WT and FSP1^TLR4LoxTB^ male or female mice (RM Two-Way ANOVA).

**Figure 3.**
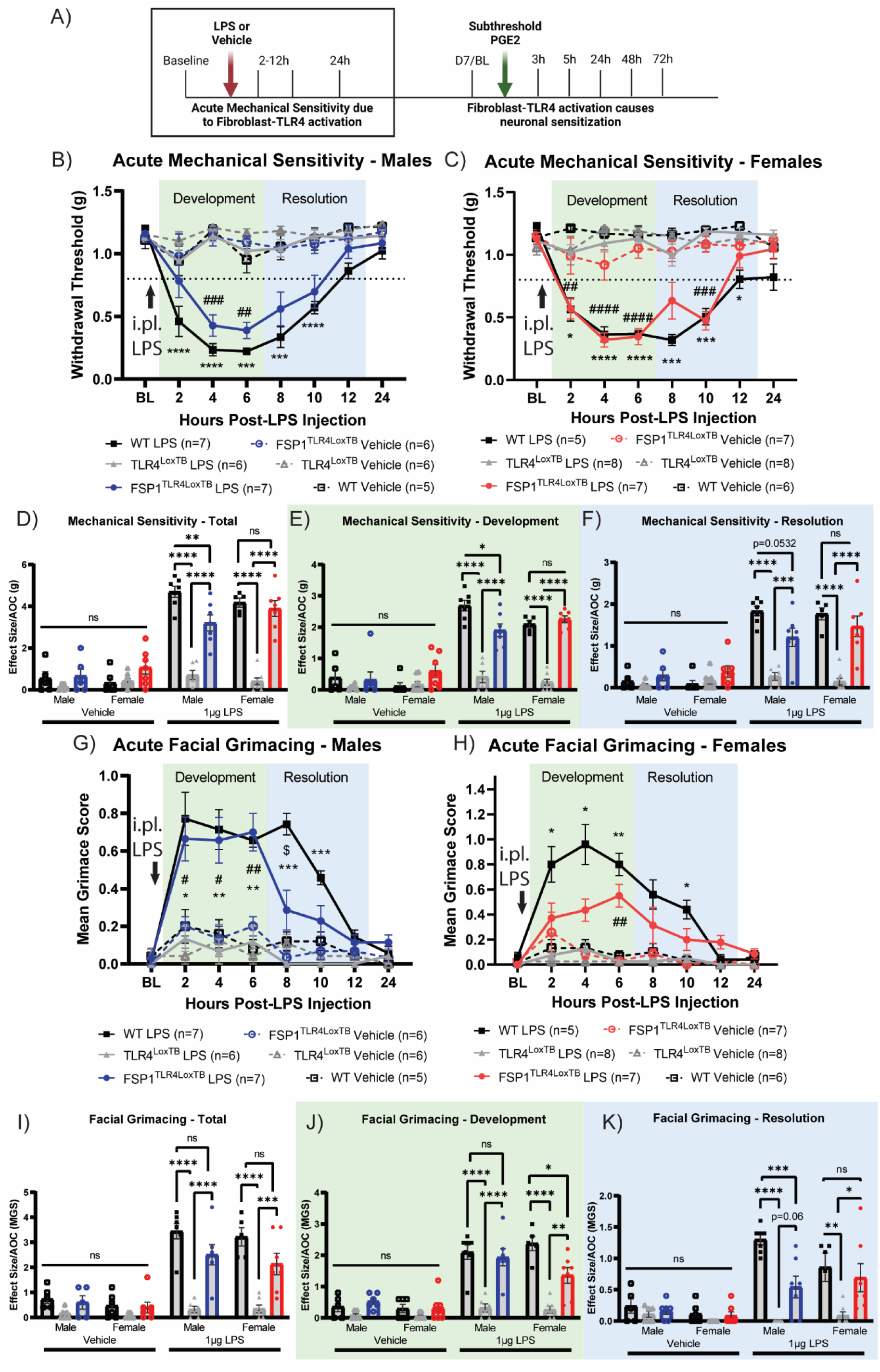
Intraplantar injection of LPS induces pain-like behavior in both male and female mice. (A) Timeline for measurement of mechanical sensitivity and facial grimacing after intraplantar injection of LPS. (B-C) Acute mechanical sensitivity of the hind paw in male (B) and female (C) mice. *WT vs TLR4^LoxTB^. #FSP1^TLR4LoxTB^ vs TLR4^LoxTB^. (D) Total effect size/AOC for mechanical sensitivity. (E) Effect size/AOC for mechanical sensitivity during the development phase (2-6h post injection). (F) Effect size/AOC for mechanical sensitivity during the resolution phase (8-12h post injection). (G-H) Acute facial grimacing in male (G) and female (H) mice. *WT vs TLR4^LoxTB^. #FSP1^TLR4LoxTB^ vs TLR4^LoxTB^. $WT vs FSP1^TLR4LoxTB^. (I) Total effect size/AOC for facial grimacing. (J) Effect size/AOC during the development phase (2-6h post injection). (K) Effect size/AOC during the resolution phase (8-12h post injection). \**p*<0.05, \*\**p*<0.01, \*\*\**p*<0.001, \*\*\*\**p*<0.0001 by Repeated Measures Two-Way ANOVA (B, C, G, H) or Ordinary Three-Way ANOVA (D-F, I-K).

Analysis of effect size (area over the curve, AOC) showed a significant reduction in overall mechanical sensitivity in FSP1^TLR4LoxTB^ males compared to their WT counterparts; However, no differences were observed between WT and FSP1^TLR4LoxTB^ female mice (**Figure 3D**, Genotype × LPS: F (2,66) = 48.62, *p*<0.0001, Three-Way ANOVA). No sex differences were observed. Effect size analysis for the development phase (2-6h post LPS) showed the same trend, with significant reduction in sensitivity in FSP1^TLR4LoxTB^ males compared to WT males and no differences between FSP1^TLR4LoxTB^ and WT females (**Figure 3E**, Genotype × LPS: F (2,67) = 37.35, *p*<0.0001, Three-Way ANOVA). During the resolution phase (8-12h post LPS), *post hoc* differences were not observed between WT and FSP1^TLR4LoxTB^ mice of either sex (**Figure 3F**, Genotype × LPS: F (2, 67) = 32.33, *p*<0.0001, Three-Way ANOVA).

We have also demonstrated that FSP1^TLR4LoxTB^ and WT mice develop spontaneous pain-like behavior as assessed via facial grimacing after intraplantar injection of LPS, whereas TLR4^LoxTB^ mice do not (**Figure 3G, H**). Although WT and FSP1^TLR4LoxTB^ males have similar development of facial grimace, FSP1^TLR4LoxTB^ males have significantly reduced grimacing at 8h post LPS compared to WT males (**Figure 3G**, F (35, 217) = 7.276, *p*<0.0001, RM Two-Way ANOVA). In contrast, FSP1^TLR4LoxTB^ females have significantly attenuated facial grimacing compared to WT females (**Figure 3H**, F (35, 245) = 6.584, *p*<0.0001, RM Two-Way ANOVA).

Analysis of effect size (AOC) showed no significant *post hoc* differences in facial grimacing between FSP1^TLR4LoxTB^ and WT male and female mice (**Figure 3I**, F (Genotype × LPS: F (2, 64) = 22.79, *p*<0.0001, Three-Way ANOVA). When assessing the development phase only, FSP1^TLR4LoxTB^ females have significantly reduced facial grimacing compared to WT females with no differences observed between male WT and FSP1^TLR4LoxTB^ mice (**Figure 3J**, F (2, 64) = 22.27, *p*<0.0001, Three-Way ANOVA). In contrast, during the resolution phase (8-12h post LPS) there was a significant reduction in facial grimacing in male FSP1^TLR4LoxTB^ mice compared to WT males with no difference observed between FSP1^TLR4LoxTB^ and WT females (**Figure 3K**, F (2, 64) = 14.66, *p*<0.0001, Three-Way ANOVA).

### Fibroblast activation leads to the development of hyperalgesic priming

To determine the effect of fibroblast activation on neuronal plasticity and transition to chronic pain, we assessed hyperalgesic priming by measuring mechanical sensitivity and facial grimacing after a subthreshold dose of PGE2 (**Figure 4A**). We administered subthreshold PGE2 (i.pl.) seven days after the resolution of LPS-induced mechanical sensitivity. Both WT and FSP1^TLR4LoxTB^ developed mechanical sensitivity after subthreshold PGE2; however, TLR4^LoxTB^ mice did not (**Figure 4B, C**).

**Figure 4.**
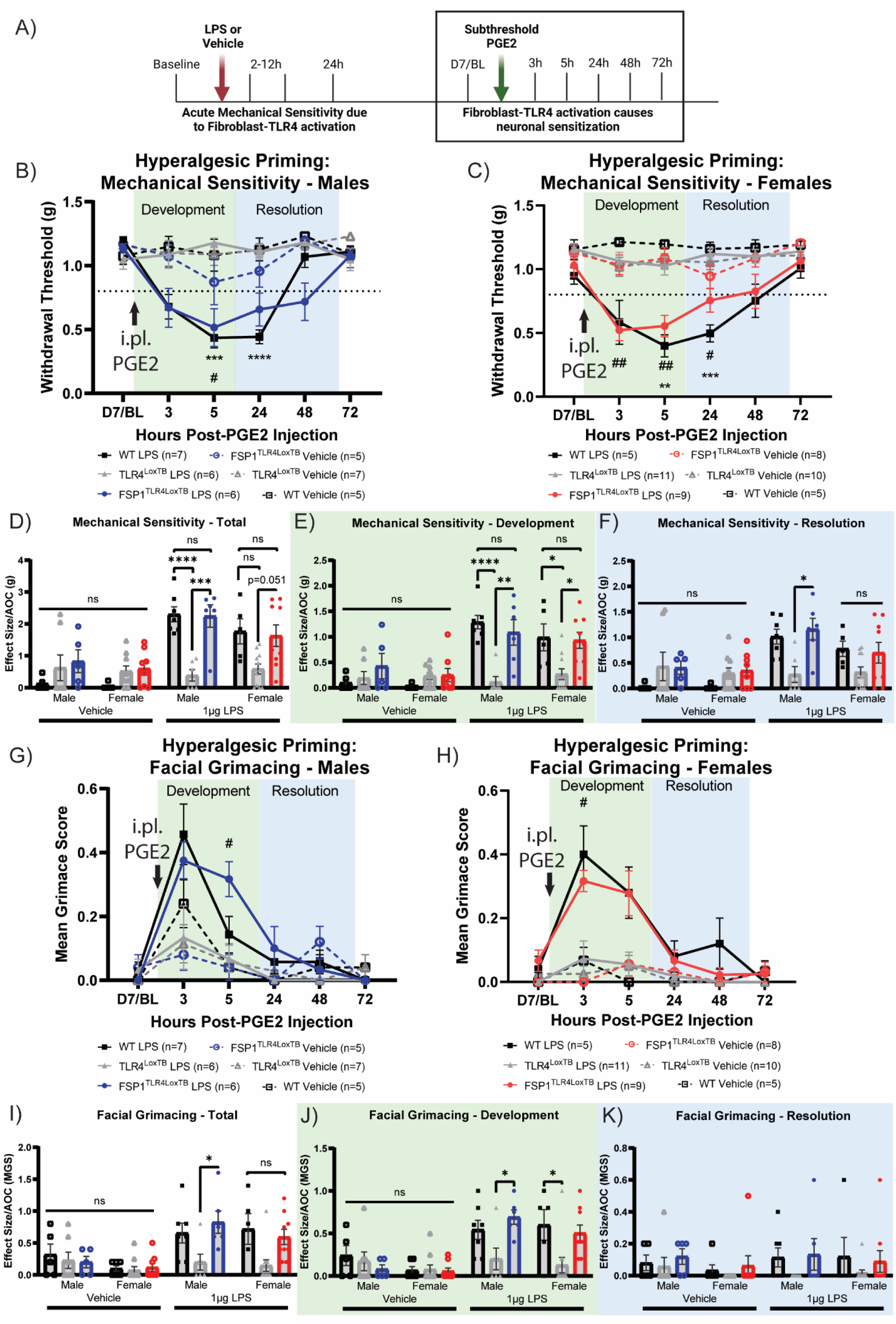
Fibroblast Activation Leads to Development of Hyperalgesic Priming. (A) Timeline for measurement of mechanical sensitivity and facial grimacing after intraplantar injection of subthreshold PGE2. (B-C) Mechanical sensitivity of the hind paw in male (B) and female (C) mice. *WT vs TLR4^LoxTB^. #FSP1^TLR4LoxTB^ vs TLR4^LoxTB^. (D) Total effect size/AOC for mechanical sensitivity. (E) Effect size/AOC for mechanical sensitivity during the development phase (3-5h post injection). (F) Effect size/AOC for mechanical sensitivity during the resolution phase (24-48h post injection). (G-H) Acute facial grimacing in male (G) and female (H) mice. #FSP1^TLR4LoxTB^ vs TLR4^LoxTB^. (I) Total effect size/AOC for facial grimacing. (J) Effect size/AOC during the development phase (3-5h post injection). (K) Effect size/AOC during the resolution phase (24-48h post injection). \**p*<0.05, \*\**p*<0.01, \*\*\**p*<0.001, \*\*\*\**p*<0.0001 by Repeated Measures Two-Way ANOVA (B, C, G, H) or Ordinary Three-Way ANOVA (D-F, I-K).

Analysis of effect size (AOC) showed that both WT and FSP1^TLR4LoxTB^ males developed significantly increased mechanical sensitivity compared to TLR4^LoxTB^ males after subthreshold PGE2 (**Figure 4D**, Genotype × LPS: F (2, 73) = 15.38, *p*<0.0001, Three- Way ANOVA). There were no differences between genotypes of female mice. During the development phase (3-5h post PGE2), both male and female WT and FSP1^TLR4LoxTB^ have significantly increased mechanical sensitivity compared to their TLR4^LoxTB^ counterparts (**Figure 4E**. Genotype × LPS: F (2, 73) = 14.51, *p*<0.0001, Three-Way ANOVA). During the resolution phase (24-48h post PGE2), only FSP1^TLR4LoxTB^ males had significantly increased mechanical sensitivity compared to their TLR4^LoxTB^ counterparts. There were no significant differences between WT mice of either sex and their TLR4^LoxTB^ counterparts, nor between FSP1^TLR4LoxTB^ and TLR4^LoxTB^ females (**Figure 4F**, Genotype × LPS: F (2, 73) = 8.793, *p*=0.0004, Three-Way ANOVA).

Male and female WT and FSP1^TLR4LoxTB^ mice develop facial grimacing after subthreshold injection of PGE2 which resolves by 24h post injection, whereas TLR4^LoxTB^ mice do not (Figure 4G, H). Interestingly, FSP1^TLR4LoxTB^ males have greater facial grimacing behavior than WT males (**Figure 4G**, F (25, 146) = 3.257, *p*<0.0001, RM Two-Way ANOVA). There were no differences observed between female WT and FSP1^TLR4LoxTB^ mice.

Analysis of effect size (AOC) showed that FSP1^TLR4LoxTB^ males exhibit facial grimacing behavior after subthreshold PGE2, with no significant effect in female mice (**Figure 4I**, Genotype × LPS: F (2, 73) = 5.952, *p*=0.004, Three-Way ANOVA). During the development phase (3-5h post PGE2), FSP1^TLR4LoxTB^ males had greater facial grimacing behavior than TLR4^LoxTB^ males, but not WT males. Conversely, only WT females had greater facial grimacing behavior than TLR4^LoxTB^ females during the development phase (**Figure 4J**, Genotype × LPS: F (2, 73) = 7.665, *p*=0.001, Three-Way ANOVA). During the resolution phase (24-48h post PGE2), no differences in facial grimacing were observed between any groups (**Figure 4K**).

### Intraplantar injection of LPS changes the morphology of dermal fibroblasts differently in male and female mice

To assess how fibroblasts change morphology in response to LPS, we injected 1µg/20µL LPS into the plantar surface of male and female FSP1^TLR4LoxTB^ mice. Hind paws were collected four hours after injection, when mechanical sensitivity was the most robust (**Figure 5A**). We used Imaris software to render 3D representations of tdTomato (^+^) cells which were analyzed for volume and cell shape characteristics (**Figure 5B**). After LPS injection, male fibroblasts are significantly smaller in volume than those from the vehicle-injected hind paw; However, no difference in volume between vehicle and LPS injected hind paws was observed in female fibroblasts (Figure 5D, G, J: LPS × sex: F (1, 16) = 24.41, *p*=0.0001, Two-Way ANOVA). No differences were observed in oblate ellipticity (flattening) in either sex (**Figure 5E**, **H, K**). After LPS injection, female fibroblasts were significantly more elongated (prolate ellipticity) than those from the vehicle-injected hind paw with no differences in males (Figure 5F, I, L: LPS × sex: F (1, 16) = 9.834, *p*=0.0064, Two-Way ANOVA).

**Figure 5.**
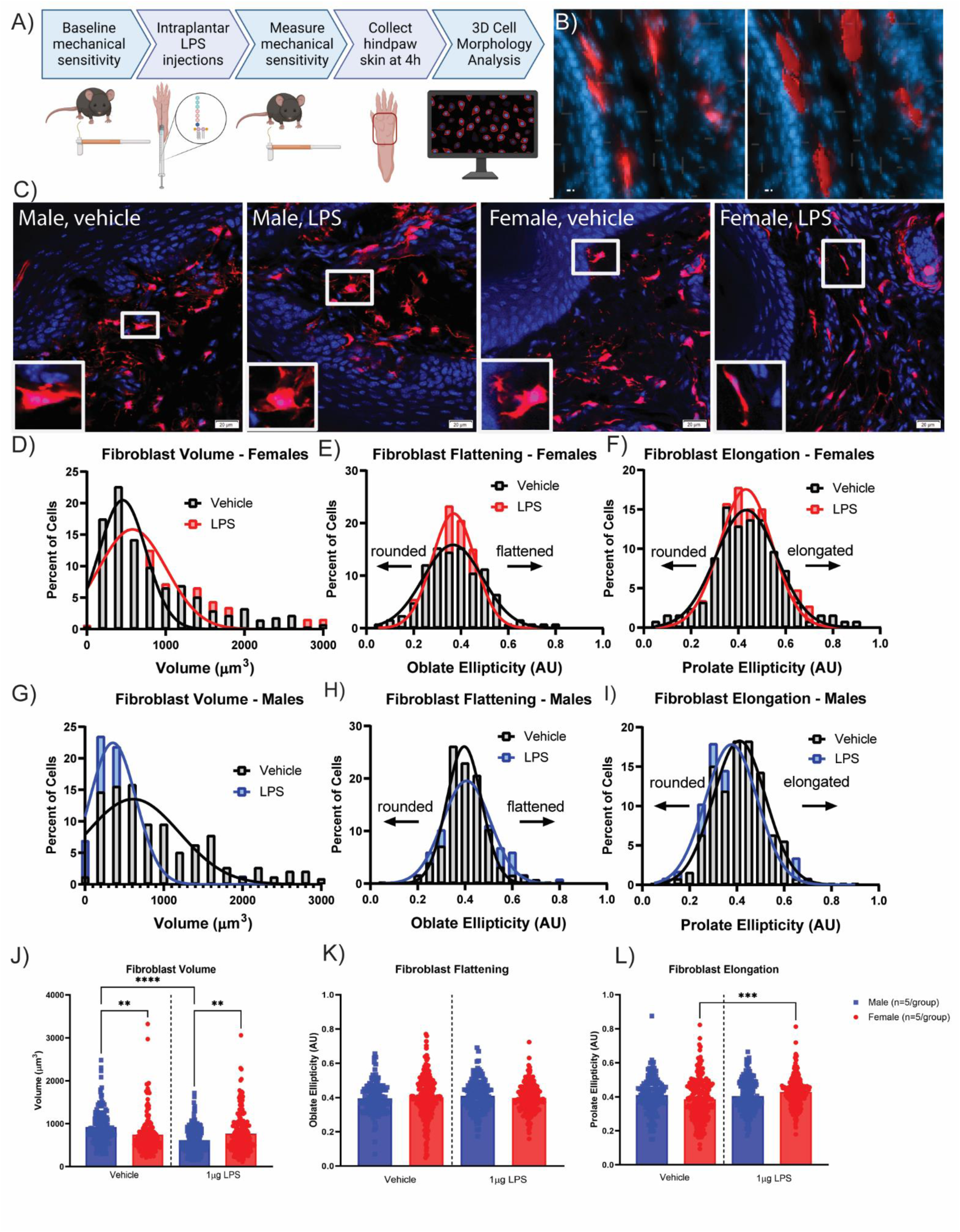
Intraplantar injection of LPS changes the morphology of dermal fibroblasts differently in males and females. (A) Experimental schematic. (B) Representative images of surface creation in Imaris software from tdTomato (^+^) cells in the hind paw dermis. Scale bar = 20µm. Original image (left) and Surfaces (right). (C) Representative confocal microscopy of tdTomato (red) and DAPI (blue) from FSP1^TLR4LoxTB^ hind paws collected 4h after intraplantar injection of LPS or vehicle. Scale bar = 20µm. Histogram of cell volume (D), cell flattening (oblate ellipticity) (E), and cell elongation (prolate ellipticity) (F) of tdTomato (^+^) dermal fibroblasts from FSP1^TLR4LoxTB^ females at 4h after intraplantar LPS or vehicle (n=5 hind paws with 70+ cells each). Histogram of cell volume (G), cell flattening (oblate ellipticity) (H), and cell elongation (prolate ellipticity) (I) of tdTomato (^+^) dermal fibroblasts from FSP1^TLR4LoxTB^ males at 4h after intraplantar LPS or vehicle (n=5 hind paws with 70+ cells each). (J-L) Fibroblast volume (J), flattening (K) and elongation (L) of male and female dermal fibroblasts at 4h after LPS or vehicle. \*\**p*<0.01, \*\*\**p*<0.001, \*\*\*\**p*<0.0001 by ordinary Two-Way ANOVA (J-L).

### Fibroblast-TLR4 activation by LPS does not acutely induce immune cells infiltration

To assess how TLR4 present on fibroblasts only affects immune response to intraplantar injection of LPS, we collected paw skin from male and female WT, TLR4^LoxTB^ and FSP1^TLR4LoxTB^ mice four hours after LPS injection at the peak of mechanical sensitivity (**Figure 6A**). Paw skin sections were stained with H&E and the number of cells in the papillary and reticular dermis of the injection site were assessed (**Figure 6B**). WT mice (regardless of sex or injection) had a greater number of cells within the papillary dermis compared to TLR4^LoxTB^ and FSP1^TLR4LoxTB^ mice (**Figure 6C**: Genotype: F (2, 12) = 7.733, *p*=0.007, Three-Way ANOVA). After LPS treatment, TLR4^LoxTB^ females had significantly reduced cell infiltration compared to WT females; However, no other differences were observed. Within the reticular dermis, no differences were observed in cell counts in vehicle injected mice; However, WT females showed significantly greater cell counts after LPS injection compared to both their vehicle-injected paw and LPS-treated TLR4^LoxTB^ females (**Figure 6D**: Genotype × LPS × Sex: F (2, 24) = 5.903, *p*=0.0082, Three-Way ANOVA). Re-expression of TLR4 in fibroblasts does not recapitulate the effects seen in WT mice of either sex. Immune cell populations within the hind paw (WT) after LPS injection were identified by CD45 and Iba1 (ionized calcium binding adaptor molecule 1) expression (**Figure 6E**).

**Figure 6.**
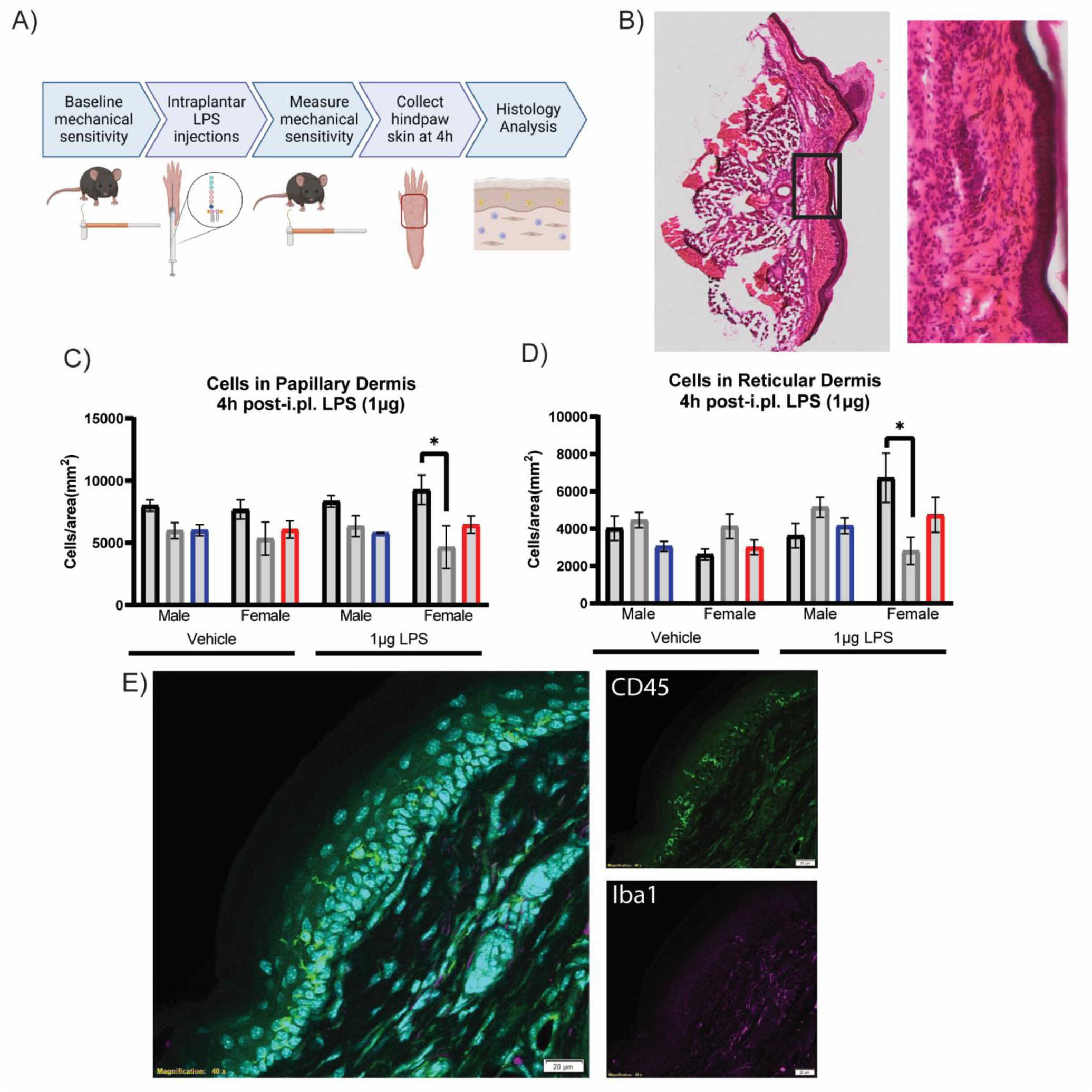
Fibroblast-TLR4 activation does not induce immune cell infiltration after LPS. (A) Experimental schematic. (B) Representative image of H&E-stained hind paw section zoomed in on the region of interest used for analysis. (C-D) Cells/mm^2^ in the papillary (C) and reticular (D) dermis at 4h post intraplantar injection of LPS or vehicle. (n=3/group). (E) Representative confocal microscopy of CD45 (green) and Iba1 (purple) in WT hind paw skin after LPS. \**p*<0.05 by Ordinary Three-Way ANOVA (C, D).

### TLR4 activation by LPS induces inflammatory cytokine production in primary dermal fibroblasts

To assess whether expression of TLR4 in FSP1^+^ fibroblasts alone was sufficient to induce cytokine production in response to LPS treatment, we measured cytokine levels in supernatant collected from cultured dermal fibroblasts (**Figure 7A**). Treatment with 1µg/mL of LPS for 24h was sufficient to induce IL-6 production in both WT and FSP1^TLR4LoxTB^ male dermal fibroblasts (**Figure 7B**: LPS treatment: (F (1.021, 13.27) = 10.76, *p* = 0.0056, Two-Way ANOVA). LPS-treated WT and FSP1^TLR4LoxTB^ fibroblasts produce significantly more IL-6 than TLR4^LoxTB^ fibroblasts as expected. Further, there was no significant difference in IL-6 production between LPS-treated WT and FSP1^TLR4LoxTB^ fibroblasts. LPS treatment also significantly upregulated IL-6 production in females (**Figure 7C**: LPS treatment: F (1.080, 21.06) = 10.52, *p* = 0.0033, Two-Way ANOVA); However, only WT females produced significantly more IL-6 than TLR4^LoxTB^ females. No significant differences were found between LPS-treated TLR4^LoxTB^ and FSP1^TLR4LoxTB^ female fibroblasts.

**Figure 7.**
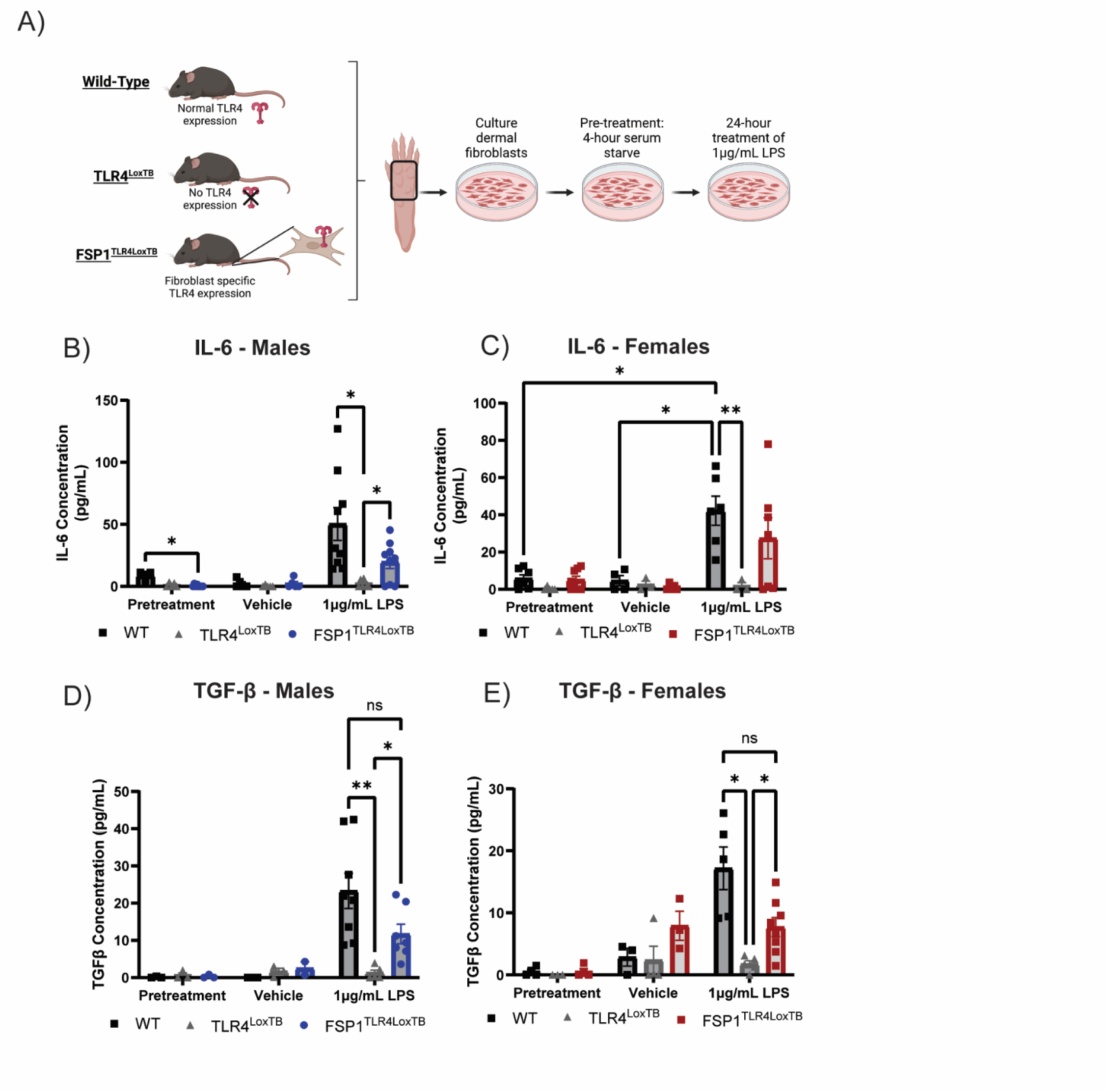
Application of LPS causes pro-inflammatory cytokine production by dermal fibroblasts. (A) Experimental timeline for treatment of cultured dermal fibroblasts with LPS. (B-C) IL-6 production by male (B) and female (C) dermal fibroblasts. (D-E) TGF-β production by male (D) and female (E) dermal fibroblasts. \**p*<0.05, \*\**p*<0.01 by Ordinary Two-Way ANOVA.

Treatment with LPS for 24h induced production of TGF-β1 in both males and female dermal fibroblasts. LPS-treated dermal fibroblasts from both WT and FSP1^TLR4LoxTB^ males produce significantly more TGF-β1 compared to TLR4^LoxTB^ male dermal fibroblasts (**Figure 7D**, F (1.019, 5.604) = 10.44, *p* = 0.0195, Two-Way ANOVA).

Further, there was no significant difference in TGF-β1 production between LPS-treated WT and FSP1^TLR4LoxTB^ male fibroblasts. Similarly, LPS-treated dermal fibroblasts from both WT and FSP1^TLR4LoxTB^ females produce significantly more TGF-β1 compared to TLR4^LoxTB^ female dermal fibroblasts (**Figure 7E**, F (1.779, 13.34) = 17.33, *p* = 0.0003, Two-Way ANOVA). There was no significant difference in TGF-β1 production between LPS-treated WT and FSP1^TLR4LoxTB^ female fibroblasts.

Treatment with LPS for 24h did not induce significant upregulation of CXCL12, IL-4, or IL-1β production in dermal fibroblasts, regardless of genotype or sex (**Table 1**).

**Table 1.** Cytokine concentrations after in vitro treatment with LPS.

|  | Pretreatment |  | Vehicle |  | 1µg/mL LPS |  |
| --- | --- | --- | --- | --- | --- | --- |
| IL-4 (pg/mL) | Male | Female | Male | Female | Male | Female |
| WT | N.D. | N.D. | 1.15 ± 1.14 | N.D. | 0.58 ± 0.58 | N.D. |
| TLR4<sup>LoxTB</sup> | N.D. | N.D. | N.D. | N.D. | N.D. | N.D. |
| FSP1<sup>TLR4LoxTB</sup> | 0.24 ± 0.12 | 0.97 ± 0.97 | 0.36 ± 0.36 | 1.87 ± 1.38 | N.D. | 1.35 ± 1.40 |
| IL-1β (pg/mL) |  |  |  |  |  |  |
| WT | 14.58 ± 2.18 | 12.21 ± 2.35 | 13.71 ± 1.95 | 11.38 ± 1.38 | 46.15 ± 5.33 | 34.45 ± 1.19 |
| TLR4<sup>LoxTB</sup> | 18.45 ± 2.26 | 8.84 ± 0.26 | 11.75 ± 1.19 | 8.80 ± 0.46 | 12.48 ± 0.30 | 15.24 ± 2.96 |
| FSP1<sup>TLR4LoxTB</sup> | 18.05 ± 4.61 | 16.09 ± 2.89 | 11.49 ± 1.55 | 9.75 ± 0.38 | 28.06 ± 2.96 | 25.35 ± 1.28 |
| CXCL12 (pg/mL) |  |  |  |  |  |  |
| WT | 36.59 ± 2.18 | 47.08 ± 7.25 |  |  | 62.64 ± 26.6 | 76.48 ± 28.1 |
| TLR4<sup>LoxTB</sup> | 35.73 ± 0.69 | 34.88 ± 0.53 |  |  | 37.84 ± 2.14 | 37.94 ± 3.06 |
| FSP1<sup>TLR4LoxTB</sup> | 38.24 ± 2.82 | 52.75 ± 16.9 |  |  | 55.99 ± 18.1 | 100.57 ± 61.8 |

Total n=3-5 per condition. N.D. = non-detectable

### FSP1^+^ cells in human skin express TLR4

To verify the translational relevance of our mouse model and current study, we performed immunohistochemistry of human skin punch biopsies to assess FSP1 and TLR4 expression in the dermis. FSP1 (+) cells can be observed throughout the dermis with similar morphology to that seen in mouse skin (**Figure** 8**A**). Furthermore, TLR4 expression can be seen in both FSP1^+^ cells and in other cell types throughout the dermis and epidermis **(Figure 8B-C)**.

**Figure 8.**
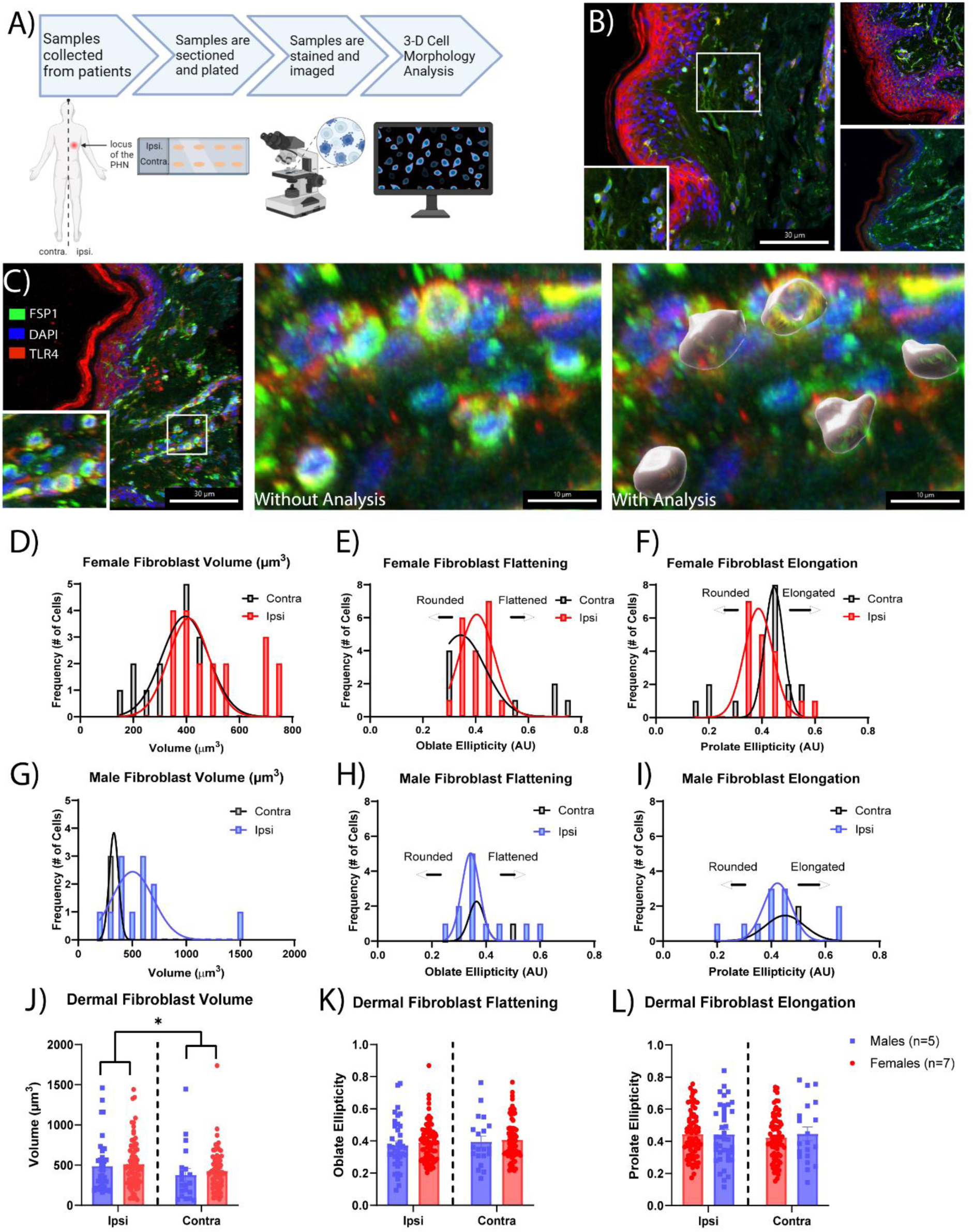
Patient skin analysis of post herpetic neuralgia dermal fibroblast morphology. (A) Experimental Schematic. (B) Representative images of patient skin stained for FSP1 (Green), DAPI (Blue), and TLR4 (Red). (C) Representative images of surface creation using IMARIS of cells positive for FSP1 overlapping with DAPI. Scale bar = 30 µm. (D) Histograms of cell volume, cell flattening (oblate), and cell elongation (prolate ellipticity). (E) Fibroblast volume, flattening, and elongation of male and female fibroblasts taken from either Ipsilateral or Contralateral skin relative to the locus of the post herpetic neuralgia neuropathy.

### Fibroblast morphology in human skin from patients with postherpetic neuralgia

To assess the translational significance of fibroblast morphology from our mouse model to human research, we analyzed human skin biopsies collected from painful dermatomes of patients afflicted by postherpetic neuralgia (PHN). We utilized painful dermatome skin from these patients as a comparison to our LPS injected mice hindpaw to further assess fibroblast morphology in a painful state. Skin biopsies were collected from sites either ipsilateral (painful) or contralateral (non-pain) dermatomes. We used Imaris software to render 3D representations of FSP1^+^ cells which were analyzed for volume and cell shape characteristics (**Figure 8C-I**). Ipsilateral dermal fibroblasts had significantly greater volume compared to contralateral fibroblasts, regardless of sex (**Figure 8E**: F (1, 207) = 4.303, p = 0.0393, Two-Way ANOVA) but no significant differences in fibroblast oblate ellipticity (flattening) or elongation (prolate ellipticity) **(Figure 8K-L)**. We found no significant *post hoc* differences in volume between male and females.

## Discussion

The current study reveals that fibroblasts are key players in pain development and hyperalgesic priming. We utilized a novel genetic animal that targets fibroblast-specific re-expression of the TLR4 gene on a whole-body null background (Jia et al., 2021). We demonstrated the functional re-expression of TLR4 on fibroblasts by showing that LPS (a potent TLR4 agonist), binds to dermal fibroblasts in mice with either wild-type expression of TLR4 (FSP1^tdt^, **Figure 1E**; WT. **Figure 2E**) or fibroblast-specific expression of TLR4 (FSP1^TLR4LoxTB^) (**Figure 2G**). Moreover, we showed the functionality of TLR4 signaling cascades as plantar injection of LPS caused pain-like behavior in male and female WT and FSP1^TLR4LoxTB^ mice, but not TLR4^LoxTB^ mice (Figure 3B-C).

Taken together, our data indicate that direct fibroblast activation mediates pain-like behavior similarly in male and female mice. In addition, fibroblast TLR4 is sufficient to induce acute mechanical hypersensitivity and hyperalgesic priming, suggesting a role for fibroblasts in the transition from acute to chronic pain. Although a few previous studies have demonstrated that fibroblasts and fibroblast-derived proteins play a role in mediating pain states (Garrity et al., 2023; Singhmar et al., 2020; Wei et al., 2014), our study is the first to show that skin fibroblast activation is sufficient to induce acute pain-like behavior and mediate the machinery involved in the transition to chronic pain.

TLR4 regulates the immune response to infection and injury via detection of pathogen-associated molecular patterns (PAMPs) and damage-associated molecular patterns (DAMPs). Fibroblasts across several tissue types utilize TLR4 signaling pathways during inflammatory states, such as uterine fibrosis, arthritis, and gingivitis, to produce cytokines and modulate the local tissue environment (Guo et al., 2015; Jian et al., 2015; Lin et al., 2015). During chronic inflammation, fibroblasts produce inflammatory factors, such as IL-6, IL-1β, and TGF-β (Garcia-Vicuna et al., 2004; Lochhead et al., 2012; Nanki et al., 2001; Wang et al., 2011). These effects are enhanced when isolated fibroblasts are treated with LPS, a potent TLR4 agonist and bacterial endotoxin (Guo et al., 2015; Lin et al., 2015; Park and Yoon, 2022). In our study, cultured WT mouse dermal fibroblasts treated with 1µg/mL of LPS produced IL-6 and TGFβ, which is consistent with previous literature (Jian et al., 2015; Lin et al., 2015; Xiao et al., 2023). These cytokines are known to activate sensory neurons to produce pain responses, thus inflammatory cytokines released by dermal fibroblasts may be part of the mechanism by which fibroblast-TLR4 activation causes acute pain responses.

Additionally, immune cell infiltration into the hind paw contributes to mechanical sensitivity after LPS (Calil et al., 2014); however, mice with fibroblast-specific expression of TLR4 did not have a significant increase in immune cell infiltration compared to either vehicle or TLR4-null mice. in . We only observed minimal production of CXCL12 by dermal fibroblasts treated with LPS, suggesting that fibroblast-TLR4 signaling does not significantly cause chemokine production during acute inflammatory responses and agonism of TLR2. Thus, our results could be due to inherent differences between dermal and synovial fibroblasts, TLR signaling mechanisms, and acute versus chronic activation of fibroblasts.

TLR4 is expressed in many cell types, including peripheral sensory neurons, myeloid-derived immune cells, endothelial cells, and more (Vaure and Liu, 2014). There are redundant TLR4-dependent pathways that are sufficient to cause pain; however, some of these pathways are both cell-specific and sex-specific. (Burton et al., 2019; Rudjito et al., 2021; Szabo-Pardi et al., 2021a). For instance, macrophage/monocyte TLR4 mediates pain states in male mice in experimental arthritis and after intraplantar HMGB1 (high mobility group box 1), whereas the presence of TLR4 on peripheral immune cells has little to no effect in female mice (Burton et al., 2019; Rudjito et al., 2021). Conversely, sensory neuron TLR4 mediates pain states in females during neuropathic pain and after intraplantar HMGB1, whereas the presence of TLR4 on sensory neurons in male mice has minimal effect on pain states (Burton et al., 2019; Szabo-Pardi et al., 2021a). In our study, we found that fibroblast TLR4 mediates both pain-like behavior and hyperalgesic priming in both males and females, suggesting that fibroblast-TLR4 signaling mediates both acute pain responses and the transition to chronic pain states in both sexes. Though the intracellular mechanisms behind this phenomenon are still being investigated, the current study provides promising evidence that targeting fibroblast TLR4 signaling pathways in therapeutic development may be an effective target for both sexes.

The relationship between cellular morphology and activation state has been demonstrated extensively in macrophages and microglia (Bertani et al., 2017; McWhorter et al., 2013). Pro-inflammatory macrophages and microglia reduce their volume and retract their processes whereas anti-inflammatory macrophages exhibit an elongated phenotype (dos Santos et al., 2023; Heinrich et al., 2017; McWhorter et al., 2013; Szabo-Pardi et al., 2021b). Fibroblast morphology is typically thought of as spindle-shaped and elongated and has primarily been studied *in vitro* using 3D cell culture systems (Lee et al., 2013; Sriram et al., 2015). “Typical” fibroblast activation (conversion from fibroblast to myofibroblast during fibrosis) includes upregulation of actin (α-SMA) and collagen (Type 1), increased cell size and a greater number of processes (Phan, 2008; Schuster et al., 2021). Therefore, we wanted to assess whether fibroblast-TLR4 activation would induce morphological changes *in vivo*. We found that fibroblasts shift morphology differently between males and females after activation via LPS-TLR4 signaling, which suggests differential intracellular signaling mechanisms.

Male dermal fibroblasts become smaller after LPS-TLR4 activation whereas female dermal fibroblasts do not change volume but become more elongated. The TLR4 activation in male dermal fibroblasts elicits a similar morphology to pro-inflammatory macrophages, suggesting shared intracellular signaling events between fibroblasts and macrophages following activation. Similar to the mouse dermal fibroblasts following intraplantar LPS injection, dermal fibroblasts in painful human PHN skin undergo changes in volume. In contrast to the male mouse dermal fibroblasts, the human fibroblasts showed an increase in volume in PHN skin whereas LPS decreased male mouse dermal fibroblast volume. We posit this difference between the mouse and human dermal fibroblast volumes may be due to the mechanism by which PHN and LPS induce pain.

Calcium dynamics in fibroblasts are thought to be an important aspect of their activation and their intra- and intercellular signaling mechanisms (Godbout et al., 2013; Rooney et al., 1989). Small, coordinated shifts in calcium oscillations are regulated by changes in second-messenger systems and intracellular calcium stores, which regulate several essential processes in fibroblasts including migration, proliferation, and collagen synthesis (Narayanan et al., 1989; Roach and Bradding, 2020; Sadras et al., 2021). Additionally, calcium signaling pathways in fibroblasts are crucial for transducing mechanical signals from their microenvironment, enabling them to respond to physical cues (Godbout et al., 2013). Furthermore, TLR4-dependent signaling pathways utilize calcium signaling mechanisms to regulate NFκB activity. Understanding and potentially manipulating these specific roles of calcium within fibroblasts offers promising avenues for therapeutic interventions aimed at regulating fibroblast activity.

Our study did not observe any potent sex differences in pain development and hyperalgesic priming after activation of fibroblast-TLR4 signaling. Interestingly, we did find some sex differences in downstream processes (morphology, calcium dynamics, and cytokine production) following the activation of TLR4 in dermal fibroblasts. In the hind paw skin of naïve mice, we saw that females had more FSP1^+^ fibroblasts compared to males (**Figure 1G**). Despite the difference in the number of fibroblasts, we observed no sex differences in the morphology of naïve dermal fibroblasts (**Supplemental Figure 1**). After LPS, female dermal fibroblasts become significantly larger and more elongated compared to males, which become smaller (**Figure 5J**).

Male and female fibroblasts may be interacting with the extracellular environment or with other cells in the dermis in different ways after activation of TLR4 signaling.

Supporting this idea, we also found that female fibroblasts consistently have greater changes to calcium oscillations after treatment with LPS compared to male fibroblasts (**Figure 1K**, **Figure 2J, Supplemental Figure 6**). While fibroblast-TLR4 signaling is sufficient to induce pain development and hyperalgesic priming in both sexes, consideration of sex is vital for understanding the mechanisms by which this occur and developing targeted therapeutics.

This study has identified TLR4 signaling on dermal fibroblasts as a signaling mechanism by which LPS causes pain-like behavior and hyperalgesic priming in both sexes. Though several studies have demonstrated that fibroblasts and fibroblast-derived proteins are involved in mediating pain states (Garrity et al., 2023; Singhmar et al., 2020; Wei et al., 2014), our study is the first to demonstrate that dermal fibroblast activation alone is sufficient to induce acute pain-like behavior and mediate the machinery involved in the transition to chronic pain. Dermal fibroblast-TLR4 activation induces several downstream processes, including morphological changes, cytokine production, and changes in intracellular calcium dynamics. However, it is unclear whether dermal fibroblasts are involved in the development and maintenance of other pain states such as post-surgical pain, neuropathic pain, or chronic pain states.

## Materials and Methods

### Animals

Adult C57BL/6 male and female mice were purchased from Jackson laboratory (stock no. 000664) and used to establish our in-house breeding colony at the University of Texas at Dallas (UT-Dallas) and used as wild-type (WT) controls for all experiments. Whole-body reactivatable TLR4-null (TLR4^LoxTB^) mice have a transcriptional blocker (LoxTB) inserted between exons 2 and 3 of the TLR4 gene and were originally a gift from Joel K. Elmquist (UT Southwestern Medical Center), but are commercially available at Jackson Laboratory (stock no. 036070) (Jia et al., 2021; Szabo-Pardi et al., 2021a). TLR4-null mice have two copies of the TLR4-LoxTB gene, have no response to TLR4 agonists, and behave as TLR4 whole-body knockouts (Jia et al., 2021). These mice have typical baseline pain measures and do not respond to TLR4 agonists such as LPS or HMGB1. These mice were crossed with FSP1cre (fibroblast-specific protein 1, or S100 calcium binding protein A4, S100A4) mice (obtained from Jackson Laboratory, stock no. 030644) to generate fibroblast specific expression of TLR4 (FSP1cre-TLR4^LoxTB^; FSP1^TLR4LoxTB^). FSP1cre miceexpress Cre recombinase under control of the *S100a4*, S100 calcium binding protein A4, promoter which is endogenously expressed in approximately 70% of stromal fibroblasts (Strutz et al., 1995; Tsutsumi et al., 2009).

All FSP1^TLR4LoxTB^ mice have two copies of the TLR4^LoxTB^ gene and a single copy of FSP1cre. The resulting mice were backcrossed for 8 generations to a C57BL/6 background at UT-Dallas. FSP1cre mice were also bred with tdTomato mice (Jackson Lab; stock no. 007909) for reporter, two-photon and immunohistochemistry experiments (FSP1cre-tdTomato; FSP1^tdt^) (**Figure 1A**). FSP1cre mice were bred with Salsa6f mice (obtained from Jackson Laboratory, stock no. 031968), which have Cre-dependent expression of tdTomato and GCaMP6f, a fluorescent calcium indicator, for calcium imaging experiments (FSP1^Salsa^) (Dong et al., 2017). Confirmation of FSP1cre- dependent expression of tdTomato was performed using the DFP Flashlight with Green LED Excitation and red filter glasses (Electron Microscopy Sciences, Cat#DFP-1) (**Figure 1B**).

Animals were kept and bred at the UT-Dallas animal facility in a pathogen free environment with standard controlled temperature (21 ± 2°C) with 12h light-dark cycle (lights on at 6:00 a.m. and off at 6:00 p.m.). Mice were housed in standard cages with 3- 5 animals per cage: standard rodent chow and water were provided *ad libitum*. Mice were 2-5 months of age for all experiments. All behavioral experiments and associated data analysis were carried out during the light cycle period. All procedures used in this study were performed in accordance with the National Institutes of Health Guidelines for the Care and Use of Laboratory Animals and in accordance with ARRIVE guidelines (National Research Council Committee for the Update of the Guide for the and Use of Laboratory, 2011; Percie du Sert et al., 2020). All experiments were carried out in accordance with protocols approved by the UT-Dallas Institutional Animals Care and Use Committee protocols 16-07 (breeding) and 17-10 (experiment).

### Polymerase chain reaction

Animals were weaned between 21-28 days of age; at which time a tail clip biopsy (∼1mm) was collected. DNA was extracted from tail clips by suspension in 75 µl extraction buffer (25 mM NaOH, 0.2 mM EDTA in ddH2O) with incubation at 95°C for one hour. An equal amount of neutralization buffer (40 mM Tris-HCl, pH 5.5) was added following incubation. DNA samples were stored at -20 °C until genotyping. PCR was performed using JumpStart REDTaq Ready Mix (Sigma, Cat#P0982-800RXN) and the primers listed below. FSP1cre was amplified using the primer pair FSPcre-FWD 5’-TCC TGC CCT TAG GTC TCA AC-3’ and FSPcre-REV 5’-CCT GTT TTG CAC GTT CAC CG-3’. TLR4 WT and LoxTB genes were amplified using the primer set TLR4TB Check F 5’-CTG ACT GGT GTG AAG TGG AAT ATC-3’, TLR4TB Check R 5’-GTC ATA GAT GCA TGC CAG ATA CA-3’, and pDisTB 5’-CTG GAC AAA CAG TGG CTG GA-3’. tdTomato WT was amplified using the primer pair oIMR9020 5’-AAG GGA GCT GCA GTG GAG TA-3’ and oIMR9021 5’-CCG AAA ATC ATC TGT GGG AAG TC-3’. tdTomato mutant was amplified using the primer pair oIMR9105 5’-CTG TTC CTG TAC GGC ATC G-3’ and oIMR9103 5’-GGC ATT AAA GCA GCG TAT CC-3’. Salsa6f WT was amplified using the primer pair 21306 5’-CTG GCT TCT GAG GAC CG-3’ and oIMR9021 5’-CCG AAA ATC TGT GGG AAG TC-3’. Salsa6f mutant was amplified using the primer pair 27656 5’-TGG TAG TGG TAG GCG AGC TG-3’ and oIMR9105 5’- CTG TTC CTG TAC GGC ATG G-3’. All primers were purchased from Integrated DNA Technologies. Samples were run on a 2% agarose (VWR, Cat# MPN605-500G) and 1X Tris-Acetate-EDTA (TAE) gel and imaged using a ChemiDoc apparatus (BioRad) and ImageLab. DNA Ladder (100 bp, Thermo Scientific GeneRuler 100 bp) was used for size determination. The FSP1cre band is at 500bp. The TLR4 WT band and TLR4^LoxTB^ bands are at 523 bp and 370 bp, respectively (**Figure 2B-C**).

### Drug delivery

#### LPS

LPS EB ultrapure (Invivogen, Cat#tlrl-pb5lps, Serotype 055:B5) was used as a potent and specific TLR4 agonist in this study, injected as 1µg LPS/20µL into the plantar surface of the hind paw using a 30G needle and Hamilton syringe (Hamilton, Cat no. 80922) (Lu et al., 2008). A 20µL injection of a 1× solution of phosphate-buffered saline (1× PBS; Research Products International Cat#P32060-04000; 10x diluted in ddH_2_O) was used as the vehicle control. Animals were observed following injections for bleeding and any other adverse response. For behavioral experiments, injections were performed between 5 and 6am, just prior to the start of the light cycle. For histology experiments, injections were performed between 10am and 11am. LPS conjugated to FITC (Sigma, Cat#F3665, Serotype 0111:B4) was used in two-photon imaging experiments to observe the uptake of LPS by cells in the dermis.

### Prostaglandin E2

To study hyperalgesic priming, a subthreshold dose of prostaglandin E2 (PGE2, Cayman Chemical Company, Cat#14010) was injected as 100ng PGE2/20µL (in 1x PBS) into the plantar surface of the hind paw using a 30G needle and Hamilton syringe (Joseph and Levine, 2010; Tierney et al., 2022). Animals were injected with PGE2 following the resolution of mechanical sensitivity after injection with LPS. There were seven days between pain resolution from LPS injections and PGE2 injections.

#### Two-Photon Imaging

To verify that dermal fibroblasts uptake and respond to TLR4 agonists *in vivo,* we used multiphoton imaging to show the direct uptake of LPS by dermal fibroblasts according to our previously published protocol (Szabo-Pardi et al., 2019). We first verified this using our reporter line, FSP1^tdt^, prior to testing this in WT, TLR4^LoxTB^, and FSP1^TLR4LoxTB^ mice. *In vivo* imaging was performed using the Olympus MPE-RS TWIN multiphoton microscope outfitted with dual excitation lasers (Spectra Physics INSIGHT DS+ -OL pulsed IR LASER, tunable from 680 to 1,300nm, 120 fs pulse width at specimen plane and SPECTRA PHYSICS MAI TAI HP DEEP SEE-OL pulsed IR LASER, tunable from 690 to 1,040nm, 100 fs pulse width at specimen plane) (Szabo-Pardi et al., 2021b).

Mice were anesthetized in the induction chamber using isoflurane followed with constant (1.5-2%) isoflurane supply using the nasal cone from the SomnoSuite Low- Flow Anesthesia System (Kent Scientific Corporation, Cat no. SS-01). Animals were then put in the prone position on the platform of the microscope using a stereotaxic apparatus. Mice were monitored throughout the experiment for any signs of distress and isoflurane concentration was adjusted accordingly. The plantar surface of hind paw was stabilized using black tape under the objective to minimize movement artifacts from respiration. Water-based lubricant was used to make an aqueous column in between the plantar skin and objective (XLPLN25XWMP2, Olympus Ultra 25x MPE water- immersion objective 1.05 NA, 2mm WD). The multiphoton microscope was manually set up using the FGR filter cube (Olympus, Cat no. FV30-FGR) to capture green and red photons from the specimen. The sapphire and Mai Tai lasers were used to excite the tdTomato (reporter tag, 1,100nm) and FITC (LPS tagged with FITC, 930nm) fluorochromes, respectively. Each hind paw was scanned using a resonant scanner with a fixed scan area of 512µm x 512µm. The dermis was imaged at 100-150 µm depth into the paw, with care taken to ensure tdTomato^+^ cells were visible to determine the correct focal plane. Baseline measurements were acquired to identify td-tomato positive fibroblasts within the dermis and ensure there was minimal autofluorescence on the FITC channel. Images were acquired over a 15-minute time lapse at 5-10 z-slices at 1µm/slice. After baseline measurements, LPS-FITC (5mg/20µL per mouse, i.pl.) was injected with care taken to not adjust the position of the hind paw. Post-injection measurements were recorded for 60 minutes using the same parameters (Szabo-Pardi et al., 2019).

#### Immunohistochemistry Reporter and Model Validation

Mice were deeply anesthetized using ketamine/xylazine and intracardially perfused with ice cold 1× PBS followed by ice cold 4% paraformaldehyde (PFA, Fisher Scientific F79). Hind paws were collected, post-fixed in 4% PFA overnight (12-16 hours), and cryoprotected in 30% sucrose (Sigma Aldrich, Cat# S0389) in 1× PBS for one week minimum. Skin from the plantar surface of the hind paw was mounted in optimal embedding/cutting medium (OCT; Thermo Fisher Scientific, Cat#50-363-773) and frozen at -80°C until sectioning. Tissue was sectioned at 20 µM thickness and mounted on positively charged microscope slides (VWR, Cat#48311-703), Sections were stored at -20°C until the immunohistochemical analysis. Frozen sections were thawed at room temperature (RT) for 15 minutes prior to tissue rehydration with 1× PBS for 5 minutes.

Tissue was pre-incubated with blocking buffer (2% normal goat serum (Gibco, Cat#16210-072), 1% Bovine Serum Albumin (VWR, Cat#97061-416), 0.05% Tween-20 (Sigma-Aldrich, Cat#P1379), 0.1% Triton X-100 (Sigma, Cat#X100) and 0.05% Sodium Azide (Sigma, Cat#RTC000068) in 1× PBS, pH 7.4) for two hours at RT to block nonspecific binding and permeabilize tissue sections. Subsequently, sections were incubated with primary antibodies (**Table 2**) diluted in blocking buffer overnight (∼20H) at 4 ⁰C. Sections were washed with 1× PBS with 0.5% Tween-20 (PBS-T) three times for five minutes each, and then incubated with secondary antibodies (**Table 2**) diluted in blocking buffer for 2H at RT. Sections were washed with 1× PBS-T solution three times for five minutes each, treated with DAPI solution for one minute (1:5000 dilution in 1x PBS-T, Sigma D9542), washed three times with 1x PBS-T, and washed once with 1× PBS. Stained sections were mounted and coverslipped using Gelvatol. Stained slides were stored at RT in the dark overnight prior to imaging.

**Table 2.** Antibodies used.

| Antibody | Company | Catalog Number | Working Dilution | RRID |
| --- | --- | --- | --- | --- |
| Anti-S100A4 mouse monoclonal IgG1 | Novus Biologicals | NBP2-53178 | 1:300 (ICC, IHC, Flow) | AB_2941095 |
| Anti-Cre Recombinase guinea pig polyclonal IgG | Synaptic Systems | 257 004 | 1:500 (IHC) | AB_2782969 |
| Anti-Collagen Type 1 rabbit polyclonal IgG | Millipore Sigma | ABT123 | 1:500 (ICC) | AB_2941296 |
| Anti-Iba1 rabbit polyclonal IgG | Fujifilm, Wako | 019-19741 | 1:500 (IHC and ICC) | AB_839504 |
| CD45 (30-411) rat monoclonal IgG2b | Thermo Fisher Scientific | 14-0451-85 | 1:500 (IHC and ICC) | AB_467252 |
| Alexa Fluor 488 goat anti-mouse IgG | Invitrogen | A11001 | 1:1000 | AB_2534069 |
| Alexa Fluor 488 goat anti-mouse IgG1 | Invitrogen | A21121 | 1:1000 (IHC and ICC); 1:500 (Flow) | AB_2535764 |
| Alexa Fluor 488 goat anti-guinea pig IgG | Invitrogen | A11073 | 1:1000 | AB_2534117 |
| Alexa-Fluor 488 goat anti-rat IgG | Invitrogen | A11006 | 1:1000 | AB_2534074 |
| Alexa Fluor 647 goat anti-mouse IgG1 | Invitrogen | A21240 | 1:1000 | AB_2535809 |
| Alexa Fluor 647 goat anti-rabbit IgG | Invitrogen | A21245 | 1:1000 | AB_2535813 |
| Alexa-Fluor 647 goat ant-mouse IgG1 | Invitrogen | A21240 | 1:1000 | AB_253580 |
| Anti-CD16/32 (purified) | eBioscience | 16-0161-85 | 1:2000 | AB_468899 |
| Anti-TLR4/MD2 Complex APC Conjugate | eBioscience | 17-9924-82 | 1:200 | AB_657858 |
| Anti-CD45 Alexa Fluor 700 Conjugate | eBioscience | 56-0451-82 | 1:200 | AB_891454 |

#### Responses to LPS

To study cellular responses to intraplantar LPS, mice were injected with 1µg LPS/20µL or 20µL of vehicle as described in section 2.2.1. Mice were injected in both the left and right hind paws with either LPS or vehicle, such that each animal acted as its own vehicle control. At four hours post-injection, mice were euthanized, and hind paws were collected and drop-fixed overnight in 4% PFA. Tissue processing, sectioning, and immunohistochemistry were performed as described above.

### Image Acquisition and Analysis

#### Reporter and Model Validation

Analysis of paw skin from FSP1^tdt^ male and female animals (n=3-4/sex, 3-5 sections/mouse) stained for Cre recombinase was performed to identify the fibroblast population in the skin that actively expresses FSP1cre. Images used for analysis were acquired using the Zeiss Axio Observer microscope equipped with an Axiocam 503 B/W monochrome camera (Carl Zeiss, Inc.). Epifluorescent z-stack images (10µm depth, 0.5µm/stack) were taken with the 20x objective (Plan-Apochromat 20x/0/80 Ph 2 M27, NA 0.8, 0.227µm/pixel) using Zen2.5 Pro software (Carl Zeiss, Inc.). Using a custom macro in ImageJ/Fiji software (Schindelin et al., 2012), the number of Cre-positive cells, the number of tdTomato-positive cells, the number of Cre-tdTomato double-positive cells, and the total number of cells in the dermis of the hind paw were assessed. Cells of interest were colocalized with DAPI. The cell population positive for both Cre and tdTomato was considered the population of interest, which was divided by the total number of cells in the dermis (via DAPI) to assess the percentage of Cre-expressing fibroblasts in the dermis. The number of cells per section was normalized via region of interest (ROI) area (µm^2^). Representative images were taken using the Olympus confocal microscope (FV1200) using Olympus Fluoview Version 4.2C software using the 40x oil objective (UPLFLN40XO NA:1.30).

### Responses to LPS

Representative images of hind paw skin immunostained for CD45 and Iba1 were taken using the Olympus confocal microscope (FV1200) using Fluoview Version 4.2C using the 40x oil objective (UPLFLN40XO NA:1.30).

### Dermal Fibroblast Cultures

Male and female mice of all genotypes were deeply anesthetized with isoflurane and euthanized via decapitation. Hind paws were thoroughly cleaned with 70% ethanol. Using a scalpel blade, skin from the plantar surface from both hind paws was removed and placed in ice-cold Hank’s Balanced Salt Solution (HBSS, divalent-free, Cat#SH30588.01). Care was taken not to collect any underlying muscle tissue. In the cell culture hood, skin tissue was removed from HBSS solution and cut into small pieces on a sterile petri dish using scalpel blades. The tissue was incubated in 1 mL of collagenase A (1mg/mL in HBSS; Sigma, Cat#10103586001) and 1 mL of collagenase D (1mg/mL in HBSS with 10% Papain (Sigma, Cat#10108014001); Sigma, Cat#11088866001) for 90 minutes in a 37°C heated water bath. The tissue was agitated via gentle shaking halfway through the incubation process. The cells were centrifuged at 1500 rcf for four minutes. The supernatant was decanted and 1 mL of trypsin inhibitor (1mg/mL enzyme and 1 mg/mL BSA in cell culture media) was added. Cell culture media used for culture experiments consists of 1× DME/F-12 1:1 (Hyclone, Cat#SH30023.01) supplemented with 2.50mM L-Glutamine, 15mM HEPES buffer, 10% Fetal Bovine Serum (FBS, HyClone Laboratories, Cat# SH3088.03), and 2% Penicillin-Streptomycin (Sigma, Cat# P4333). The tissue was triturated using filtered P1000 pipette tips until the solution became cloudy. The solution was filtered through a 70-µm nylon cell strainer (Corning, Cat#431751) and washed with an additional 9mL of culture media. The suspension was centrifuged at 1500 rcf for four minutes and the cell pellet was resuspended in 1 mL of media for counting. Fibroblasts were plated at 100,000 cells/mL and maintained in culture for 5-7 days. After the first media change (24h post plating), the penicillin/streptomycin concentration was reduced to 1% for all subsequent media changes and treatments.

### LPS Treatment & Immunocytochemistry

Fibroblasts were plated on poly D-lysine (PDL, 2µg/mL, Sigma-Aldrich, Cat# P0899- 10MG) coated coverslips (Corning BioCoat 12mm, Fisher Scientific, Cat# 08-774-384) at a density of 100,000 cells/mL in a 24-well cell culture plate. Fibroblasts were maintained for 5-7 days until 90% confluency. Cells were incubated in serum-free media for four hours, and then treated with LPS (1µg/mL in serum-free media) for 24 hours.

Supernatant was collected and immediately frozen at -80°C until use for measurement of cytokine levels (described below). Cells were washed with 1x PBS three times and fixed with pre-chilled 4% PFA for 15 minutes. Cells were washed three times with 1x PBS and incubated in blocking buffer for two hours at room temperature. Cells were then incubated in primary antibody cocktail, diluted in blocking buffer, overnight (∼20 hours) at 4°C. Cells were washed three times with 1X PBS-T and incubated in secondary antibody cocktail, diluted in blocking buffer, for two hours at RT and protected from light. Cells were washed three times with 1X PBS-T, incubated with DAPI (1:5000) for one minute, washed twice with 1X PBS-T, then washed once with 1X PBS. Coverslips were mounted on uncharged microscope slides using Gelvatol. After mounting, slides were stored in the dark at RT overnight prior to imaging.

Representative images were taken using the Olympus confocal microscope (FV1200) using Olympus Fluoview Version 4.2C software using the 40x oil objective (UPLFLN40XO NA:1.30).

### Cytokine Measurement via ELISA

Supernatant from LPS-treated fibroblast cultures was assayed for the following cytokines: IL-6 (Mouse IL-6 DuoSet ELISA kit, R&D Systems, Cat# DY406-05), IL-4 (Mouse IL-4 ELISA MAX Deluxe Set, BioLegend, Cat#431104), IL-1β (Mouse IL-1β/IL- 1F2 DuoSet ELISA kit, R&D Systems, Cat#DY401), TGF-β (Legend Max Mouse Free Active TGF-β1 ELISA kit, BioLegend, Cat#437707) and chemokine: CXCL12/SDF-1β (Legend Max Mouse CXCL12 (SDF-1β) ELISA kit, BioLegend, Cat#444207). All assays were performed according to manufacturer’s instructions. All samples were assayed in duplicate. All ELISAs were performed by experimenters blinded to condition.

### Calcium Imaging – Fura-2 AM

For calcium imaging experiments, fibroblasts from WT, TLR4^LoxTB^, and FSP1^TLR4LoxTB^ mice were plated on 35mm glass-bottom dishes (MatTek, Cat#P35G-1.5-10-C) with No. 1.5 Coverslip and 10mm diameter at 100,000 cells in 100µL cell culture media. All dishes were coated with PDL prior to plating. Plated cells rested in the cell culture incubator for two hours prior to flooding with cell culture media. Cells were maintained for 5-7 days prior to calcium imaging experiments until reaching 80% confluency.

On the day of imaging, each dish (WT, TLR4^LoxTB^, and FSP1^TLR4LoxTB^ mice) was loaded with 10µL fura-2 AM (Invitrogen, #F1221) in 1mL loading buffer (1x HBSS containing 0.25% w/v BSA (endotoxin-free; Sigma-Aldrich, Cat#A9576) and 2mM CaCl_2_) for one hour at 37°C. After the fura-2 AM incubation, plates were incubated in 2mL normal bath solution (125 mM NaCl (Fisher Scientific, Cat#S271-500), 5mM KCl (Fisher Scientific, Cat#P217-500), 10mM HEPES (Sigma-Aldrich, Cat#H4034), 1M CaCl_2_ (Sigma-Aldrich, Cat#21115), 1M MgCl_2_ (Fisher Scientific, Cat#M35-500), and 2M glucose (Sigma-Aldrich, Cat#G7528) in ddH2O) at pH of 7.4 ± 0.05 and mOsm of 300 ± 5 for 30 minutes at 37°C.

All experiments were performed using the MetaFluor Fluorescence Ratio Imaging Software on an Olympus TH4-100 apparatus using a 40x oil objective. For experiments using fura-2 AM, ratiometric changes of bound (excitation: 340nm) to unbound (excitation: 380nm) fura-2 AM (emission: 510nm) were recorded in real time (1 frame/second). Baseline recordings were measured for 60 seconds, during which time normal bath solution was applied to cells. Normal bath supplemented with LPS (1µg/mL) was applied for 300 seconds followed by a 120 second wash with normal bath, and finally a 30 second application of 100µM ATP (made in normal bath, Sigma Aldrich, Cat#A6419) as a positive control. High concentrations of ATP have been shown to elicit robust and rapid influx of calcium in several cell types (Lembong et al., 2015; Zumerle et al., 2019).

### Calcium Imaging - FSP^Salsa^

Experiments using FSPSalsa mice were performed as described above with the following adjustments. Fibroblasts from FSP^Salsa^ mice were plated on uncoated or collagen-coated (MatTek, Cat#P35GCOL-1.5-10-C) dishes. All dishes were coated with PDL prior to plating. Fibroblasts from each mouse were plated on two PDL-coated dishes and two collagen-coated dishes.

The day prior to imaging, two dishes from each mouse (one PDL and one collagen) were changed to serum-free culture media and incubated at 37°C overnight. All other dishes received a normal media change. On the day of imaging, fibroblasts were incubated in normal bath solution for 30 minutes at 37°C prior to imaging. Regions of interest were first selected based on expression of tdTomato to confirm FSP1cre expression and then GCaMP6 intensity changes were recorded in real time (1 frame/second) using the FITC channel.

Previous studies have demonstrated that cellular calcium responses are comparable when using fura-2 AM and GCaMP based calcium imaging (Kim et al., 2014; Tierney et al., 2022). We have opted to keep these datasets separated to examine differences between general fibroblast calcium activity and FSP1^+^ fibroblast calcium activity.

### Calcium Oscillation Analysis

Data from each plate was exported to an associated excel sheet that plotted the change in fluorescence intensity of each cell at one-second intervals. As a positive control, each cell was screened for an ATP response (>15% increase in fluorescence intensity) (Lembong et al., 2015). For experiments using fura-2 AM, cells with baseline fura-2 AM ratios (F_340nm_/F_380nm_) less than 0.45 or greater than 1.75 were excluded. The data was further split into 3 sections based on treatment: Baseline (bath) (0-60 seconds), LPS (60-360 seconds), and post-LPS (bath) (360-480 seconds). Each data set was then run through a data analysis program written in python using pandas (1.3.5), NumPy (1.21.5), Matplot libraries (3.2.2), and SciPy(1.4.1) and run using Google Colab (version 2023/06/23) to measure oscillations per minute in the cells. Excel files (.xlxs files, raw fluorescence data) was imported using pandas. The fluorescence intensity data was then inputted into the program which removed all the points nonlocal minima or maxima using the find_peaks() function from SciPy to speed up processing time later on.

The peaks were found by the find_peaks(), function from SciPy, the data was multiplied by -1 then fed into find_peaks() to find troughs. Both troughs and peaks were appended back together to create a data set with only local maxima and minima in order using .append(). The corresponding times of these maxima/minima points were put into a separate array. This condensed data set was then run through a separate find_peaks() using prominence. Prominence uses relative heights to aid in noise extraction. find_peaks() uses horizontal lines on a digital elevation model to find intersections of other peaks; peaks of the same elevations are ignored. Using the peak information, find_peaks() finds the minimum values on each side, this is the peak’s bases. The higher base is the lowest contour line of the peak. Prominence is the height difference between the height of the peak and its base. This prominence was done for all local maxima. The prominence was set to the Interquartile range (IQR) of the data multiplied by a constant of 0.4, which captured the largest number of peak values, above noise. These prominent peaks were then counted to determine how many peaks were in the data. A prominent peak to peak would quantify 1 cycle/oscillation. The data was then graphed as oscillations per minute using the Matplotlib plotting functions, marking all of the prominent peaks **(Supplemental Figure 1E-F**). This process was repeated for every cell in the set. A text and excel file were generated after the data was processed. Details about specific functions utilized can be found in **Table 3**.

**Table 3.**

| Function | Library | Description |
| --- | --- | --- |
| np.percentile | numpy | Calculates the given percentile of an array. |
| find_peaks() | scipy.signal | Identifies peaks in a signal. |
| plt.text() | matplotlib.pyplot | Adds text at specified coordinates on a plot. |
| plt.xlabel() | matplotlib.pyplot | Sets the plot label for the x-axis. |
| plt.ylabel() | matplotlib.pyplot | Sets the plot label for the y-axis. |
| plt.title() | matplotlib.pyplot | Sets the plot title. |
| plt.plot() | matplotlib.pyplot | Plots the given data. |
| plt.savefig() | matplotlib.pyplot | Saves the current figure to a file in various formats. (.PNG) |
| pd.read_csv() | pandas | Reads a comma-separated values (.CSV) file into a DataFrame. |
| pd.DataFrame() | pandas | Creates a DataFrame object. |

Data are represented as the number of oscillations per minute (oscillation frequency) for each cell. Cells were excluded if they had a calcium oscillation frequency of less than 1 oscillation per minute during either baseline or LPS recordings. For FSP^Salsa^ mice, no female fibroblasts were excluded due to lack of calcium oscillations; however, 3.5% of male FSP^Salsa^ fibroblasts were excluded. Additionally, one calcium imaging dish (male FSP^Salsa^ in a PDL-coated dish) was not used in final data analysis due to lack of calcium oscillations in >90% of cells in the dish. All calcium imaging data was analyzed by independent experimenters blinded to condition.

### Flow Cytometry

Plantar skin from male and female mice (WT, TLR4^LoxTB^, and FSP1^TLR4LoxTB^) was collected and dissociated as described in section 2.5 with the following modifications. Flow buffer (1x dPBS + 0.5% endotoxin-free BSA and 0.02% glucose) was used to wash cells through 70-micron strainers and final cell suspensions were in flow buffer. To block non-specific binding via the Fc receptor, cells were incubated in blocking buffer (anti-CD16/32 purified antibody in flow buffer) for 20 minutes on ice. Samples were incubated with pre-conjugated extracellular flow antibodies (TLR4, CD45) diluted in flow buffer for 60 minutes, on ice and protected from light. Prior to intracellular staining, cells were carefully washed with flow buffer and incubated with BD Fixation/Permeabilization solution (Fisher Scientific, Cat#BDB554715) for 20 minutes on ice, protected from light. Cells were carefully washed with 1x BD Perm/Wash Buffer (diluted from 10x in ddH_2_O) and incubated with intracellular antibody (FSP1/S100A4 mIgG1) diluted in 1x BD Perm/Wash Buffer for 45 minutes, on ice and protected from light. Cells were carefully washed with 1x BD Perm/Wash Buffer and incubated with secondary antibody (Goat anti-mIgG1 Alexa Fluor 488) diluted in 1x BD Perm/Wash Buffer for 45 minutes, on ice and protected from light. Samples were carefully washed with 1x BD Perm/Wash buffer and resuspended in flow buffer for data acquisition. Appropriate compensation controls, isotype controls, and secondary controls were used for determination and gating.

Stained samples were analyzed using a BDAria Fusion and FlowJo software. Experimenters were blinded to genotype. Antibody details can be found in **Table 2**.

### Behavioral experiments

All behavioral measurements were assessed during the light cycle. Animals were handled for a minimum of 5 minutes each prior to acclimation to the experimental room. Mice were then acclimated to the experimental room while in their home cages for a minimum of one hour. Mice were habituated to the behavior racks for 4h for minimum 2 days before data acquisition. Baseline measures were conducted on two separate days prior to injection. Mice were randomly assigned to receive drug or vehicle by an independent experimenter. On testing days, mice were acclimated in acrylic behavior boxes for approximately one hour prior to testing. Spontaneous pain (grimace scale) and mechanical allodynia (Von Frey) assessments were conducted in the von Frey racks that were 11 cm long by 10 cm wide and 4.5 in height. Behavior racks were cleaned with a 1:3 ratio of deodorant-free cleaner (Seventh Generation^TM^, 22719BK-5) to eliminate odor cues. Mice were given time in their home cages with free access to rodent chow and water between the 4h and 6h, and then between the 8h and 10h time points for behavior. Experimenters were blinded to the genotype and treatment group.

### Facial Grimacing

Spontaneous pain was assessed by observation of facial expression using the Mouse Grimace Scale (Langford et al., 2010; Lenert et al., 2022). Five components of facial grimacing were assessed: orbital tightening, ear position, cheek bulge, nose bulge, and whisker position. Each component was scored as follows: not present was given a score of ‘0’, moderately present was given a score of ‘1’, and obviously present was given a score of ‘2’. The score of the five components was averaged to give a mean grimace score (MGS). Spontaneous pain measures were assessed prior to evoked pain measures at each timepoint.

### von Frey test

Animals were tested for mechanical hypersensitivity using the von Frey assay (Chaplan et al., 1994; Lenert et al., 2022). Paw withdrawal thresholds were assessed in response to the application of calibrated von Frey hair filaments using the up-down method.

Filaments with logarithmically incremental stiffness of 2.83, 3.22, 3.61, 3.84, 4.08, 4.17 (converted to the 0.07, 0.16, 0.4, 0.6, 1.0, 1.4 g, respectively) were applied to the plantar surface of the hind paw. To avoid tissue damage, a cut off of 2g was applied. A positive response was noted by paw withdrawal, licking, or shaking of the paw. The withdrawal of the ipsilateral and contralateral paws was measured and recorded separately.

### Fibroblast Morphology *in vivo*

To assess the effect of activation via LPS-TLR4 on 3D *in vivo* fibroblast morphology, FSP1^TLR4LoxTB^ mice were given an intraplantar injection of 1mg/20mL of LPS or vehicle (1× PBS). Each animal was given two injections: one LPS injection and one vehicle injection in opposite hind paws. The drug injections were randomized to either the left or right hind paw for each animal. Hind paw collection and processing was performed as described in section 2.3.2. Frozen sections were thawed at room temperature (RT) for 15 minutes prior to tissue rehydration with 1× PBS for 5 minutes. incubated in DAPI (1:5000) for one minute, washed with 1× PBS, and mounted with Gelvatol. Slides were stored in the dark at RT overnight prior to imaging.

Images for analysis were acquired using the Zeiss Axio Observer Microscope equipped with an Axiocam 503 B/W monochrome camera (Carl Zeiss, Jena, Germany) using the z-stack mode of the ZEN software for DAPI and tdTomato expression. Epifluorescent z-stack images (10µm depth, 0.5µm depth) were taken with 20x objective (NA 0.8) 0.227µm/pixel using ZEN2.5 Pro software (Carl Zeiss, Inc,). Images were taken at 14-bit depth as .czi files. Three images and a minimum of 70 cells were analyzed per hind paw.

Image analysis was performed using Imaris software (Oxford Instruments, version 9.0.1). Z-stack images are uploaded into Imaris where they are reconstructed into voxels and create a 3D object that spans across the z-stack. Image files (.czi files) are uploaded into Imaris’s Arena (.ims files) and the 3D reconstruction was viewed using the Surpass function. The Surfaces function was used to create 3D objects using background subtraction for tdTomato positive cells within the dermis, which were cross referenced with DAPI. Surface creation parameters were based on the most representative images and were made using the Imaris Surfaces creation wizard. Two filters based on object size and tdTomato fluorescence intensity were used to remove artifacts. Only tdTomato positive cells with a visible nucleus (assessed using DAPI) were considered for this analysis. The volume, sphericity, and ellipticity of tdTomato positive cells in the dermis was assessed. The sphericity of an object is given as a value from 0.01-1.00 with 1.00 being a perfect sphere and the closer to zero being more ellipsoid. Prolate ellipticity is given as a value from 0.01-1.00, with values closer to zero being more spherical and values closer to 1 being more ellipsoid. An object with high prolate ellipticity has one axis significantly longer than the other two and can be considered elongated. Oblate ellipticity is given as a value of 0.01-1.00, with values closer to zero being more spherical and values closer to 1 being more ellipsoid. An object with high oblate ellipticity has two axes similar in length but longer than the third axis and is a more flattened shape (Szabo-Pardi et al., 2021b, **Supplemental Figure 1A)**. Analysis was performed by experimenters blinded to the sex and injection condition. Representative images for publication were acquired using the Olympus confocal microscope (FV1200) using Olympus Fluoview Version 4.2C software using the 40x oil objective (UPLFLN40XO NA:1.30).

#### Mouse Paw Skin Histology (H&E)

To assess the infiltration of cells into the paw skin following intraplantar injection of LPS, mice were given an intraplantar injection of 1µg LPS/20µL or 20µL vehicle (1× PBS) as described above. Paw skin was collected, processed, and sectioned as described above. Sections were stained with Hematoxylin (Sigma, Cat. no HHS16) and Eosin Y Solution (Sigma-Aldrich, Cat. no 318096). Stained sections were imaged via brightfield microscopy using the Olympus VS120 Virtual Slide Microscope at 40x magnification.

Analysis of 4-8 serial sections per paw was performed using the Olympus CellSens Software. Cell counts for the papillary and reticular dermis of the paw sections were focused on the injection site. Analysis was performed by experimenters blinded to the sex, genotype, and injection.

#### RNA in situ hybridization - Mouse

We performed RNA *in situ* hybridization on fresh frozen DRGs (L3-L4) and hind paw skin from WT mice to assess *tlr4* and *s100A4* mRNA localization. RNAscope *in situ* hybridization assay (Multiplex Fluorescent Reagent Kit version 2) was performed following the Advanced Cell Diagnostics (ACD) protocol using the following adjustments. Samples were incubated in fresh 10% neutral buffered formalin (NBF) at 4 °C for one hour and a 30-second Protease IV digestion was used for all experiments. The fluorophores used were from Akoya Biosciences at a 1:1500 dilution: TSA Plus Fluorescein (Cat# NEL741001KT) for C1 and TSA Plus Cyanine 5 (Cat# NEL745001KT) for C3. We used the following probes: Mm-S100a4 (Cat#412971) and Mm-Tlr4-C2 (Cat#316801-C2). The RNAscope 3-plex negative control probe (Cat#320871), which targets the bacterial gene DapB, was used to confirm probe specificity and control for background staining. To check for RNA and tissue quality, the RNAscope 3-plex positive control probe- Mm (Cat#320881) was used to target the genes UBC (high expressing), PPIB (moderate-high expressing), and POLR2A (moderate-low expressing). Images were taken using the Olympus confocal microscope (FV1200) using Olympus Fluoview Version 4.2C software using the 40x oil objective (UPLFLN40XO NA:1.30).

### Human skin biopsy immunohistochemistry and morphology

#### Human skin immunohistochemistry

Human skin punch biopsies (provided by Integrated Tissue Dynamics (INTiDYN; Renesselaer, NY) were obtained from painful and non-painful dermatomes of post- herpetic neuralgia (PHN) patients participating in a non-related phase 2 clinical trial (registered on clinicaltrials.gov as NCT02365636; patient and experimental details are described elsewhere (Fetell et al., 2023)). Briefly, skin punch biopsies were collected from painful, PHN-affected dermatomes (Ipsilateral) and unafflicted dermatomes (Contralateral). Biopsies were cryoprotected, sectioned at 14µm thickness perpendicular to the skin surface, and mounted together across a series of individual slides, such that each slide contains individual patient tissue sections from ipsilateral and contralateral biopsies. Tissue slides were maintained under glycerol/PBS at -80°C until use.

Frozen tissue section slides were thawed at room temperature (RT) for 15 minutes prior to tissue rehydration with 1× PBS for 5 minutes. Tissue sections were incubated with blocking buffer (2% normal goat serum (Gibco, Cat#16210-072), 1% Bovine Serum Albumin (VWR, Cat#97061-416), 0.05% Tween-20 (Sigma-Aldrich, Cat#P1379), 0.1% Triton X-100 (Sigma, Cat#X100) and 0.05% Sodium Azide (Sigma, Cat#RTC000068) in 1× PBS, pH 7.4) for two hours at RT to block nonspecific binding and permeabilize tissue sections. Subsequently, sections were incubated with primary antibodies (**Table 2**) diluted in blocking buffer overnight (∼20H) at 4 ⁰C. Sections were washed with 1× PBS with 0.5% Tween-20 (PBS-T) three times for five minutes each, and then incubated in the corresponding secondary antibody cocktail diluted in blocking buffer for 2 hours at RT. Sections were washed with 1× PBS-T solution three times for five minutes each, treated with DAPI solution for one minute (1:5000 dilution in 1x PBS-T, Sigma D9542), washed three times with 1x PBS-T, and washed once with 1× PBS. Immunolabled sections were mounted and coverslipped using Fluoroshield. Labeled slides were stored at RT in the dark overnight prior to imaging.

Images were taken using the Olympus confocal microscope (FV1200) using Olympus Fluoview Version 4.2C software using the 60x oil objective (UPLSAPO60XO NA:1.35).

### Acquisition and analysis

Image acquisition and analysis was performed by experimenters blind to sex, age, treatment, and location of the PHN neuropathy. Images for analysis were acquired on the Olympus confocal microscope (FV1200) using Olympus Fluoview version 4.2C software using the 40x oil objective (UPLFLN40XO NA:1.3). Images were taken at an objective of 40x with z-spacing of approximately 1µm. Images were exported as a (.oib) filetype to be imported into Imaris 10.0.1. The image analysis was performed in Imaris (Oxford Instruments, version 10.0.1). Imaris automatically converts .oib files to its native filetype (.ims). After conversion, 3-D reconstructions from the z-stacks taken on the Olympus confocal were automatically re-compiled via the built-in software in Imaris.

Surface creation, a method within Imaris to generate rendered shapes based on parametric criteria, was used to generate the outlines of the fibroblasts. A border was created to separate the dermis from the epidermis. Three criteria were used in the analysis: visible cell nuclei (assessed via DAPI), dermal location, and if the shape of the cell could be completed. The last criteria prevented cells which were along the edge of the image from being counted as their morphologies would be incomplete.

Additionally, this also prevented overlapping cells from being counted as one cell, rather than two. The first surface generated was for DAPI using surface creation and setting the channel to DAPI. A morphological split filter was used in the surface creation, allowing Imaris to generate a linked surface rather than one surface. The second surface identified FSP1^+^ cells by calibrating the surface creation tool to the FSP1^+^ channel (488), and the morphological split filter was applied. In the creation of the FSP1 surface, two filters were added to ignore all surfaces which did not contain overlap with DAPI and to disable surface creation within the epidermal layer. This allowed for analysis of dermal FSP1^+^ fibroblasts only. The classifications of volume, sphericity, and ellipticity of the FSP1^+^ cells were collected from the data tab under specific values in the Imaris program.

### Morphological Classification

The volume, sphericity, and ellipticity of FSP1^+^ cells in the dermis were assessed. The sphericity of an object is given as a value from 0.01-1.00 with 1.00 being a perfect sphere and the closer to zero being more ellipsoid. Prolate ellipticity is given as a value from 0.01-1.00, with values closer to zero being more spherical and values closer to 1 being more ellipsoid. An object with high prolate ellipticity has one axis significantly longer than the other two and can be considered elongated. Oblate ellipticity is given as a value of 0.01-1.00, with values closer to zero being more spherical and values closer to 1 being more ellipsoid. An object with high oblate ellipticity has two axes similar in length but longer than the third axis and is a more flattened shape (Szabo-Pardi et al., 2021b). Image acquisition and analysis was performed by experimenters blind to sex, age, treatment, and location of the PHN neuropathy.

Representative images for publication were acquired using the Olympus confocal microscope (FV1200) using Olympus Fluoview Version 4.2C software using the 40x oil objective (UPLFLN40XO NA:1.30).

### Statistical analysis

Statistical analysis was performed using GraphPad Prism software (Version 10.0.2). All datasets were expressed as mean ± standard error (SEM). Statistical significance for all tests was set at *p* < 0.05. Comparison of two groups was performed using unpaired two-tailed t-test, whereas comparison of three or more groups was performed using ordinary one-way ANOVA. Behavioral data (raw data) was analyzed via Repeated Measured Two-Way ANOVA with *post hoc* Tukey’s multiple comparisons test. Behavioral data (effect size/area over the curve (AOC)) was analyzed via Ordinary Three-Way ANOVA with *post hoc* Tukey’s multiple comparisons test. For morphology datasets, histograms were made by calculating the frequency distribution for each dataset and plotting the percentage of values within each bin. Sex and treatment comparisons were performed using Ordinary Two-Way ANOVA with *post hoc* Tukey’s multiple comparisons test. Cell counts for H&E-stained paw skin were analyzed using Ordinary Three-Way ANOVA with *post hoc* Tukey’s multiple comparisons test. Cytokine data was analyzed using Ordinary Two-Way ANOVA with *post hoc* Tukey’s multiple comparisons test.

## Supplemental Material Summary

### Data Availability

The data are available from the corresponding author upon reasonable request.

## Acknowledgements

The authors would like to thank Brandon T Lane for his technical assistance for animal behavior experiments and Dalton E Moore for his technical assistance and training with Imaris software. We would also like to thank the current and past members of the NIB Lab for their contributions to this work. This work was supported by NIH grant F99NS129173 (MEL), Eugene McDermott Graduate Fellowship 202205 (MEL), R21DK130015-01A1 (MDB), Rita Foundation Award in Pain (MDB), and the University of Texas Rising STARS program research support grant (MDB).

## CRediT authorship contribution statement

Melissa E Lenert: Conceptualization, Data Curation, Formal Analysis, Investigation, Methodology, Writing – Original Draft and Review and Editing

Nilesh M Agalave: Conceptualization, Data Curation, Formal Analysis, Investigation, Methodology, Writing – Original Draft

Emily K. Debner: Data Curation, Investigation, Formal Analysis, Writing – Review and Editing

Syed A. Naqvi: Formal Analysis, Software, Writing - Review and Editing Andreas M. Chavez: Formal Analysis, Software, Writing - Review and Editing Jessica A. Tierney: Investigation, Methodology, Writing – Review and Editing Marilyn Dockum: Resources, Methodology, Writing – Review and Editing Phil Albrecht: Resources, Writing – Review and Editing

Frank Rice: Resources, Writing – Review and Editing Theodore J. Price: Resources, Writing – Review and Editing

Erica L. Sanchez: Methodology, Writing – Review and Editing, Resources

Michael D. Burton: Conceptualization, Data Curation, Formal Analysis, Funding Acquisition, Investigation, Methodology, Resources, Supervision, Writing – Original Draft and Review and Editing

## Abbreviations

AOC: area over the curve
DAMPs: damage-associated molecular patterns
DRG: dorsal root ganglia
ECM: extracellular matrix
FB: fibroblast
FSP1: fibroblast-specific protein 1
HMGB1: high mobility group box 1
Iba1: ionized calcium binding adaptor molecule 1
MGS: mean grimace score
NBF: neutral buffered formalin
OCT: optimal embedding/cutting medium
Osc: oscillations
Osc/min: oscillations per minute
PAMPs: pathogen-associated molecular patterns
PBS-T: 1x PBS + 0.05% Tween-20
PDL: poly-D lysine
PFA: paraformaldehyde
PGE2: prostaglandin E2
PRR: pattern-recognition receptor
RM: repeated measures
ROI: region of interest
RT: room temperature
S100A4: S100 calcium binding protein A4

